# Nucleus Pulposus Cell Network Modelling in the Intervertebral Disc

**DOI:** 10.1101/2024.09.18.613636

**Authors:** Sofia Tseranidou, Maria Segarra-Queralt, Francis Kiptengwer Chemorion, Christine Le Maitre, Janet Piñero, Jérôme Noailly

## Abstract

Intervertebral disc degeneration (IDD) arises from an intricate imbalance between the anabolic and catabolic processes governing the extracellular matrix (ECM) within the disc. Biochemical processes are complex, redundant and feedback-looped, thus improved integration of knowledge is needed. To address this, a literature-based regulatory network model (RNM) for nucleus pulposus cells (NPC) is proposed, representing the normal state of the intervertebral disc (IVD) cells, in which proteins are represented by nodes that interact with each other through activation and/or inhibition edges. This model includes 32 different proteins and 150 edges by incorporating critical biochemical interactions in IVD regulation, tested *in vivo* or *in vitro* in human and animal NPC, alongside non-tissue-specific protein-protein interactions. We used the network to calculate the dynamic regulation of each node through a semi-quantitative method. The basal steady state successfully represented the activity of a normal NPC, and the model was assessed through the published literature, by replicating two independent experimental studies in human normal NPC. Pro-catabolic or pro-anabolic shifts of the network activated by nodal perturbations could be predicted. Sensitivity analysis underscored the significant influence of transforming growth factor beta (TGF-β) and interleukin-1 receptor antagonist (IL-1Ra) on the regulation of structural proteins and degrading enzymes within the system. Given the ongoing challenge of elucidating the mechanisms that drive ECM degradation in IDD, this unique IVD RNM holds promise as a tool for exploring and predicting IDD progression, shedding light on IVD phenotypes and guiding experimental research efforts.

## Introduction

In 2020, the global prevalence of LBP exceeded half a billion cases. This condition was responsible for 7.7% of all Years Lived with Disability (YLDs), making it the leading cause of disability worldwide^1^. IDD is one of the main unique causes of LBP, as it explains about 26%-42% of cases^2^. The IVD is the largest avascular organ in the human body, located between the vertebral bodies and consisting of three main tissues: the nucleus pulposus (NP); the annulus fibrosus (AF); and the cartilage endplate (CEP). Due to the avascular nature of the organ, the nutrient supply to NPC is made through diffusion over the interstitial fluid of the ECM, resulting in a slow tissue turnover. IDD is multifactorial with a number of inter-relating risk factors including genetic inheritance, age, metabolic factors, mechanical loads, and other environmental factors, altogether leading to altered cellular phenotypes. These alterations translate into a disruption of the balance between the anabolic and catabolic processes that regulate the ECM of the disc, tilting the balance towards catabolic pathways (see Fig.1), ultimately contributing to the onset of LBP.

**Figure 1.**
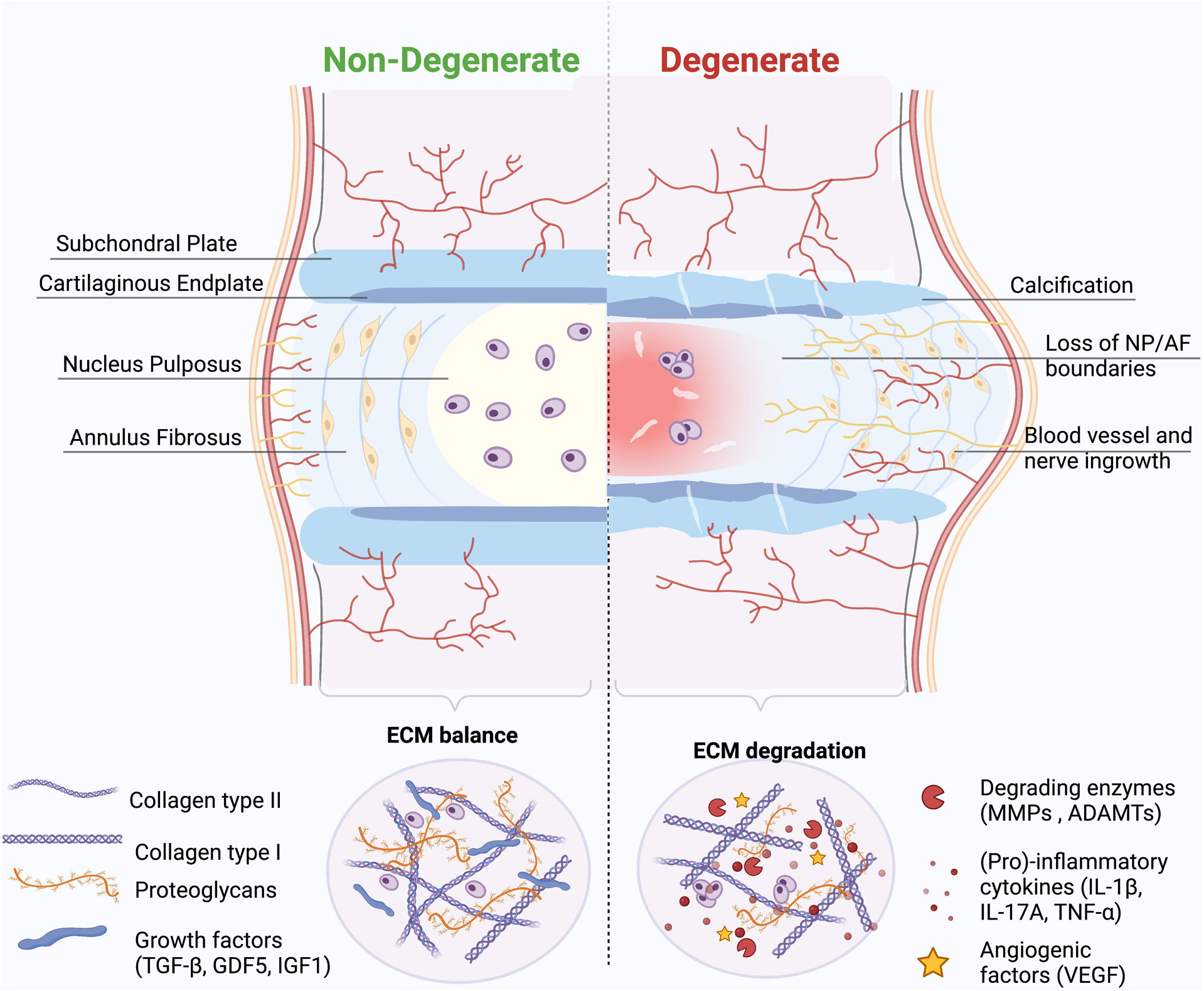
Intervertebral Disc. Structure and main ECM components of a non-degenerate, IVD (Left) and a degenerate IVD (right). Within a non-degenerate human IVD the anabolic and catabolic components that regulate the ECM of the disc are kept in balance. During disc degeneration the balance is dysregulated, resulting in decreasing matrix synthesis of COL2A and ACAN and promotion of degrading enzymes (MMPs, ADAMTs). Regarding the disc morphology clear boundaries between NP and AF are difficult to distinguish with degeneration. Created with BioRender.com (2024).

**Figure 2.**
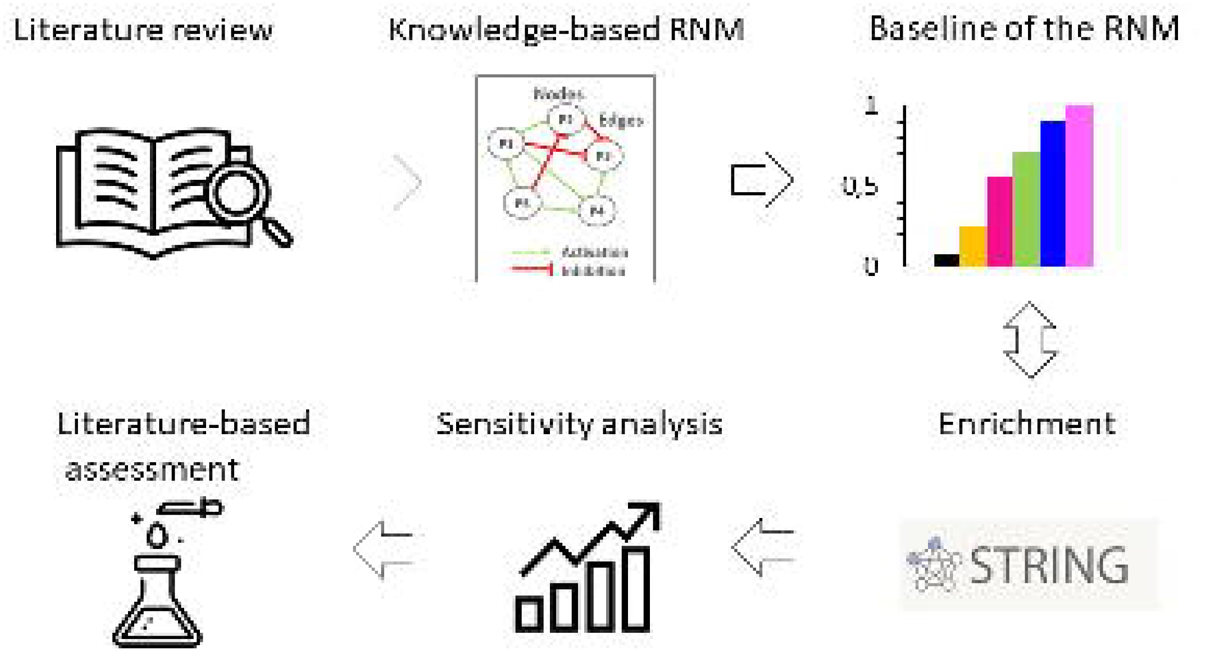
Overview workflow. A literature review was firstly conducted to collect all the information about biochemical stimulation and activity of NPC in IVD regulation. Then, a knowledge-based RNM was built, and its baseline was calculated and analysed. The following enrichment method fed continuously the network with new connections from general protein-protein interactions in Homo Sapiens. The RNM was re-assessed, and a sensitivity analysis identified the cytokines and growth factors with the highest impact in the system, which was again compared to independent literature knowledge.

During disc degeneration the native NP and AF cells of the IVD can acquire pro-inflammatory phenotypes^3^ that might trigger pro-catabolic events with a strong inertia over time. The early stage of degeneration involves a cascade of cellular changes resulting in loss of proteoglycans and thus associated hydration, and type-II collagen, in the NP.

Several experimental studies have been conducted to explore the biochemical environment of the IVD, which is responsible for the onset and the progression of its degeneration. Some have focused on the catabolic effect of pro-inflammatory cytokines on the ECM^4–7^ and how the inhibition of these cytokines could help in IVD treatment^8–10^. Other studies have explored the role of anti-inflammatory cytokines, such as IL-4 and IL-10^11–13^, and the potential role of growth factors, including GDF5^14^ in the alleviation of IDD through cell proliferation and differentiation, reduced cell apoptosis, and tissue regeneration.

Despite the avail of anti-inflammatory and pro-anabolic factors, it is difficult, however, to understand the initiating factors (disc microenvironment, biomechanics, genetics and epigenetics, bacterial infection of the disc and the gut microbiome) involved in the shift from anabolism to catabolism, by using only *in-vivo* or *in-vitro* techniques ^15^. Computational methods, like Finite Element Models (FEM), offer valuable insights by simulating the mechanical loading of the IVD structure in both healthy and degenerated states. These models can predict changes in disc tissue structure, composition, and properties, including cellular activities such as nutrition^16–19^. However, FEM require the use of constitutive models and laws with well-defined quantitative parameter, posing challenges when modelling targets at cellular and/or sub-cellular levels. Given the necessity of capturing regulatory mechanisms at these finer scales to understand IDD, modelling approaches must grapple with the intricacies of these interactions, often relying on knowledge or semi-qualitative biological measurements.

According to such a need, biological networks offer graph-based knowledge representations in a holistic and intuitive way. In these models, proteins are depicted as nodes interacting through activation and/or inhibition links. Static interaction maps are transformed into computable models to predict cellular responses, either through logical relationships among nodes and links, or via ordinary differential equations (ODEs). When knowledge needs to be mapped to cope with the lack of specific laws, ODE can be derived through Boolean rules or fuzzy-logic interpolations thereof ^20,21^. The latter can provide information about relative (or pseudo levels of) protein concentrations or expression, on a continuous normalized scale, and effectively recognize key cell regulators responsible for the ECM disruption. Indeed, numerous studies employing cell network models have aimed to pinpoint key molecules in IDD, with potential applications in diagnosis, disease monitoring, and therapeutic interventions ^22–25^.

Xu et al. (2021) provided fresh insights into the regulatory network of IDD, by integrating proteomics and transcriptomic profiling data from degenerated human NP cells^22^. Their analysis unveiled six key regulators (TNFAIP6, CHI3L1, KRT19, DPT, COL6A2 and COL11A2) with potential functional roles in the IDD process, and they identified two regulators already reported (KRT19, COL6A2) as important IDD markers in independent studies. Li et al. (2022) conducted a comprehensive genome-wide study on RNA expression profiles, constructing protein and disease-gene interaction networks, thereby identifying three novel IDD-specific genes^23^. Moreover, they identified entrectinib and larotrectinib as potential treatments for IDD via the NTRK2 gene^23^. Huang et al. (2021) constructed a regulatory mechanism network of lncRNA/circRNA– miRNA–mRNA in IDD ^24^. They proposed potential interaction axes, such as NEAT1–miR-5100–COL10A1 and miR663AHG/HEIH/hsa-circ-0003600–miR-4741–HAS2/HYAL1/LYVE1, shedding light on molecular mechanisms in IDD, notably implicating type X collagen synthesis and hyaluronic acid metabolism imbalance in degeneration. Finally, a top-down network modelling approach was proposed to estimate NPC responses in multifactorial environments ^21,26^, by systematically translating experimental data into feed-forward model parameters, facilitating high-level simulations of cell activity based on factors such as cell nutrition and pro-inflammatory activity.

Despite notable progress, there remains a lack of models to capture the regulation of NPC activity or phenotype, which is complex due to the multitude of pro-anabolic and pro-catabolic factors, which impact on the complex biochemical environments within the NP in non-degenerate IVDs and during IDD. Within articular cartilage, Segarra-Queralt et al. (2022) has developed a knowledge-driven network, with a verified balance between pro-anabolic and pro-catabolic processes, which was calibrated and validated against relative protein concentrations^27^. However, to date no such model has been developed to simulate the complex interactions and regulation within the IVD.

Hence, this study aims to develop an extensive knowledge driven network model to simulate IVD cell activity, augmenting current approaches in IVD systems biology. Our approach prioritizes high-level modelling, focusing solely on the effects of proposed interactions among soluble proteins, without delving into the intricacies of intracellular signalling mechanisms.

This facilitates assessments against falsifying experiments and retain the bare essential descriptors of normal and perturbed cell activity, while preserving model interpretability^27^. Furthermore, the present network modelling approach provides a curated corpus, shared hereby, that uniquely reflects the activity of NPC in terms of regulated soluble factors, according to current knowledge about non-degenerate IVD and IDD.

## Results

### Overview

An initial literature-based network model was meticulously crafted, capturing the complex anabolic and catabolic protein interactions within NPC, critical for ECM regulation in the IVD. After first simulation assessments, this model underwent enrichment by integrating general protein-protein interactions sourced from the Homo sapiens dataset within the String database, along with additional information about closely related cells such as chondrocytes. Following this enrichment process, the model successfully portrayed the anticipated NPC baseline behaviour within a non-degenerate IVD. To further validate its qualitative accuracy, the model underwent thorough testing against independent experimental data concerning the activity of human non-degenerate IVD NPC in *in vitro* cell culture treatments that favoured either anabolic or catabolic responses. Moreover, a sensitivity analysis was conducted through a full factorial design of experiments to pinpoint the soluble regulators exerting the most significant impact on structural proteins and degrading enzymes within the network. Specifically, emphasis was placed on identifying cytokine and growth factor nodes with the greatest influence. In essence, this approach yielded a unique and knowledge-driven NPC model capable of replicating expected cellular activity within the context of a non-degenerate IVD. Both the model and the underlying literature corpus were shared.

### Knowledge-based RNM

From the literature review, we gathered information about soluble proteins that were shown to affect NPC activity. The initial topology consisted of 31 different nodes (proteins) and 59 edges that represented node interactions in terms of activations or inhibitions (Fig.3). Full network detail can be found in the following repository^28^. Pro-inflammatory cytokines and degrading enzymes had the highest degree of connectivity (DoC), with IL-1β being dominant with 28 DoC. Conversely, only few connections linked the anti-inflammatory/anti-catabolic and pro-anabolic regulators such as IL-10 (2 DoC), IL-4 (5 DoC), TGF-β (1 DoC), IGF (1 DoC), GDF5 (4 DoC) and TIMP (2 DoC), to the rest of the network.

**Figure 3.**
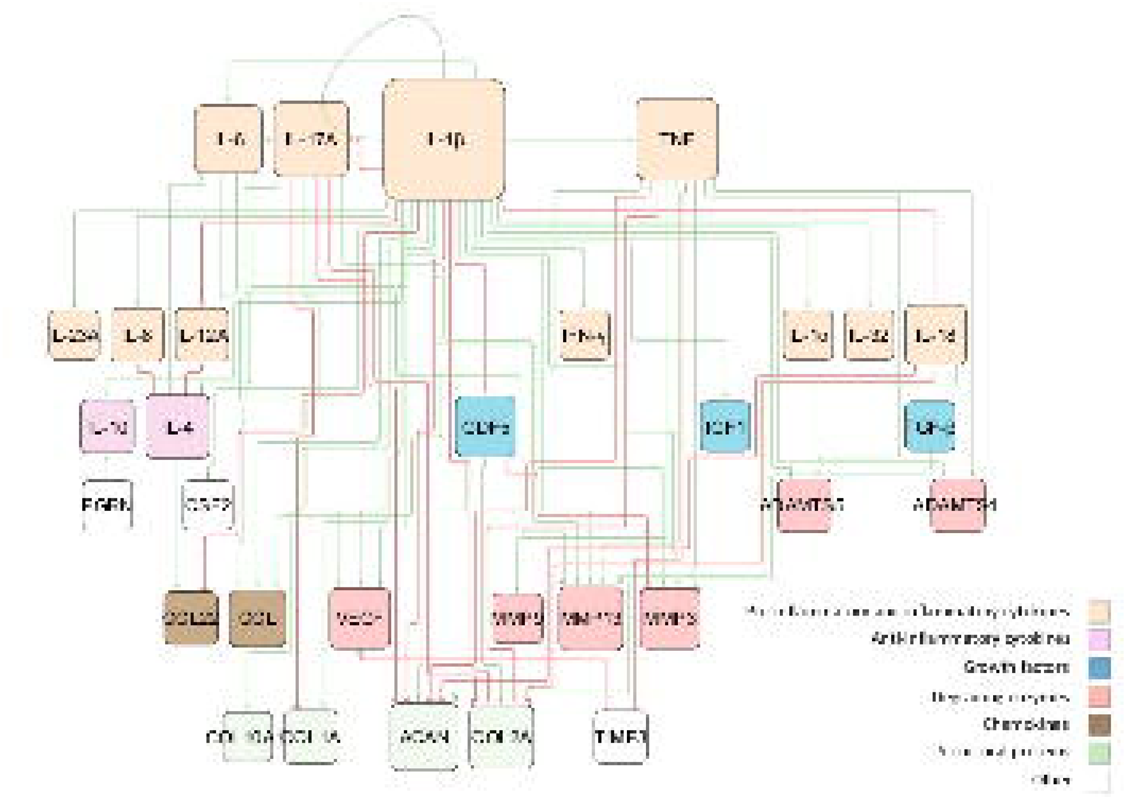
Topology of the initial RNM. The network consists of high-level protein-protein interactions, including the most important soluble proteins in IVD regulation. These are connected through activation (green lines) and inhibition (red lines) links. The size of the node is proportional to the number of connections, i.e. nodes with bigger size have more connectivity.

Node activations were computed by using a semi-quantitative determination of the SS of the network, detailed elsewhere^20^. The exact code used can be found in the corresponding repository^29^ while the model was deposited in BioModels^30^ and assigned the identifier MODEL2407080001. After conducting 100 simulations to ensure stability, we examined the distribution of the resulting SS for each protein. These distributions were skewed, with small variance of the SS values. To enhance clarity for readers, we chose, therefore, to represent the median of the baseline using bar plots (see Fig. 4), while the detailed boxplot analyses are available in the Supplementary Figure 1. The activation of pro-catabolic or abnormal matrix components such as COL10A1, COL1A, IFN-γ, TNF, IL-1α, IL-1β, IL-6, CCL, MMP3, MMP9, MMP13, ADAMTS4/5 and VEGF was notably abundant. Conversely, only a few anabolic / anti-catabolic components, including IL-4, IL-10, IGF1 and GDF5 were observed to be activated. Interestingly, the activation levels of IVD extracellular matrix proteins nodes, COL2A and ACAN, were calculated to be very low.

**Figure 4.**
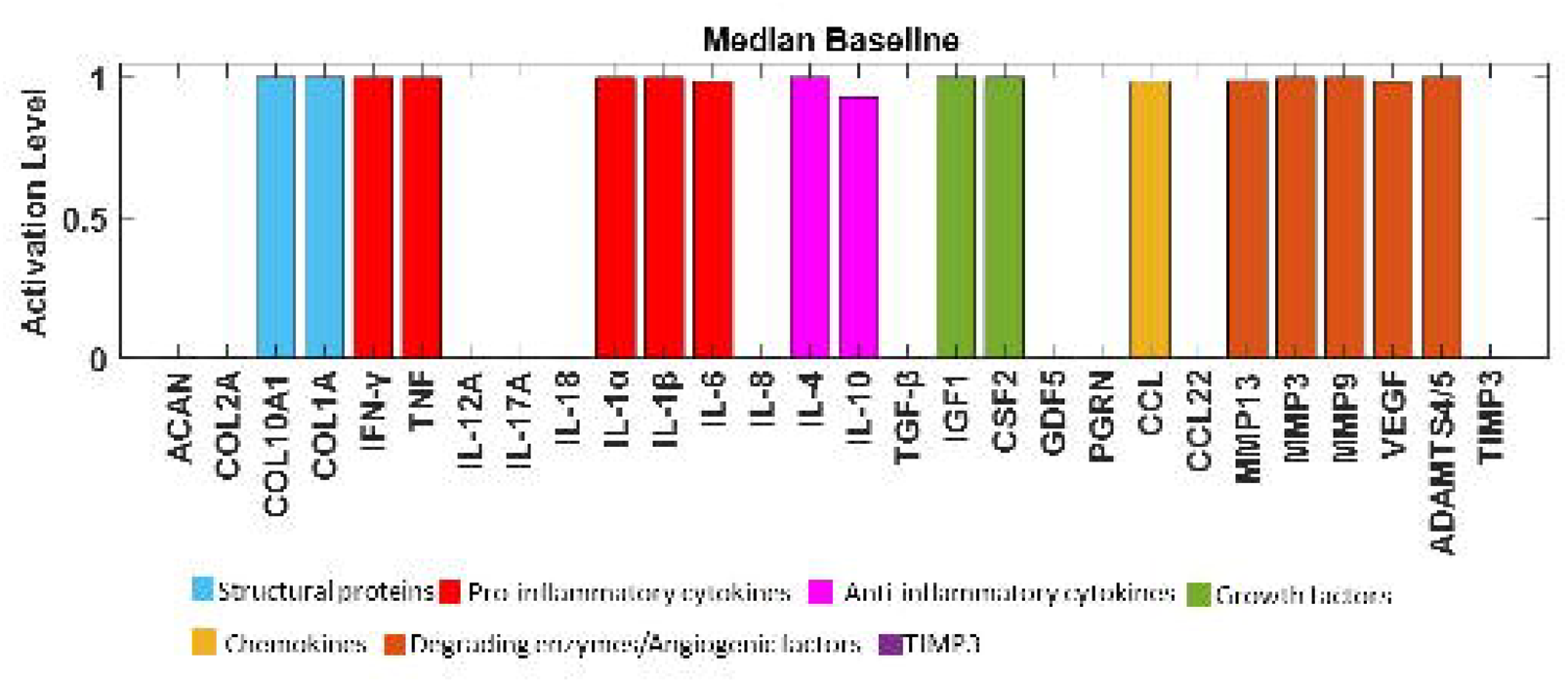
Median baseline of the initial RNM. The baseline of the knowledge-based regulatory network is represented through bar plots and reflects a degenerate state of the disc. Activation level (0 for no activation and 1 for the highest activation), of each regulator is given in the SS of the system and are colour-labelled depending on their biochemical properties in the IVD regulation.

### Enriched RN model

The first objective was to build a regulatory network that reaches a SS representative of a non-degenerate IVD, i.e., with a SS characterized by high pro-anabolic node activity and low pro-catabolic activations. However, the initial topology converged to a clearly pro-catabolic SS (Fig.3). This motivated the enrichment process through the STRING database (Szklarczyk et al. Nucleic Acids Res. 2015 43(Database issue): D447-52), aimed at transitioning the network from representing a pro-catabolic state to a pro-anabolic state of the IVD. As a result, general protein-protein interactions in Homo sapiens, along with relevant interactions in chondrocyte regulation, were incorporated. The topology of the enriched network (found in the corresponding repository^28^), highlighted the importance of the potential missing links, including anti-inflammatory cytokines such as IL-10 (18 DoC) and IL-4 (19 DoC), growth factors like TGF-β (15 DoC), IGF1 (8 DoC), and GDF5 (6 DoC), as well as inhibitors of matrix degradation: TIMP1/2 (9 DoC) and TIMP3 (3 DoC) (Fig.4). The enriched model included 32 different proteins and 150 interactions. Compared to the initial RNM, we included four new proteins (Fig.5, denoted by bold borders) and 91 new links (Fig.5, represented by bold lines). Some commonly known or expected interactions such as IL-4 inhibits IL-1β, IL-10 inhibits IL-1β, TGF-β inhibits ADAMTS4/5 and TGF-β inhibits MMP2 could not be found in the IVD-specific literature but were reflected in the STRING database. The updated enriched network was finally able to represent a pro-anabolic activity with high expression level of structural proteins and growth factors and low expression level of (pro)-inflammatory cytokines and degrading enzymes (Fig.6). The detailed boxplot analyses are available in the Supplementary Figure 2.

**Figure 5.**
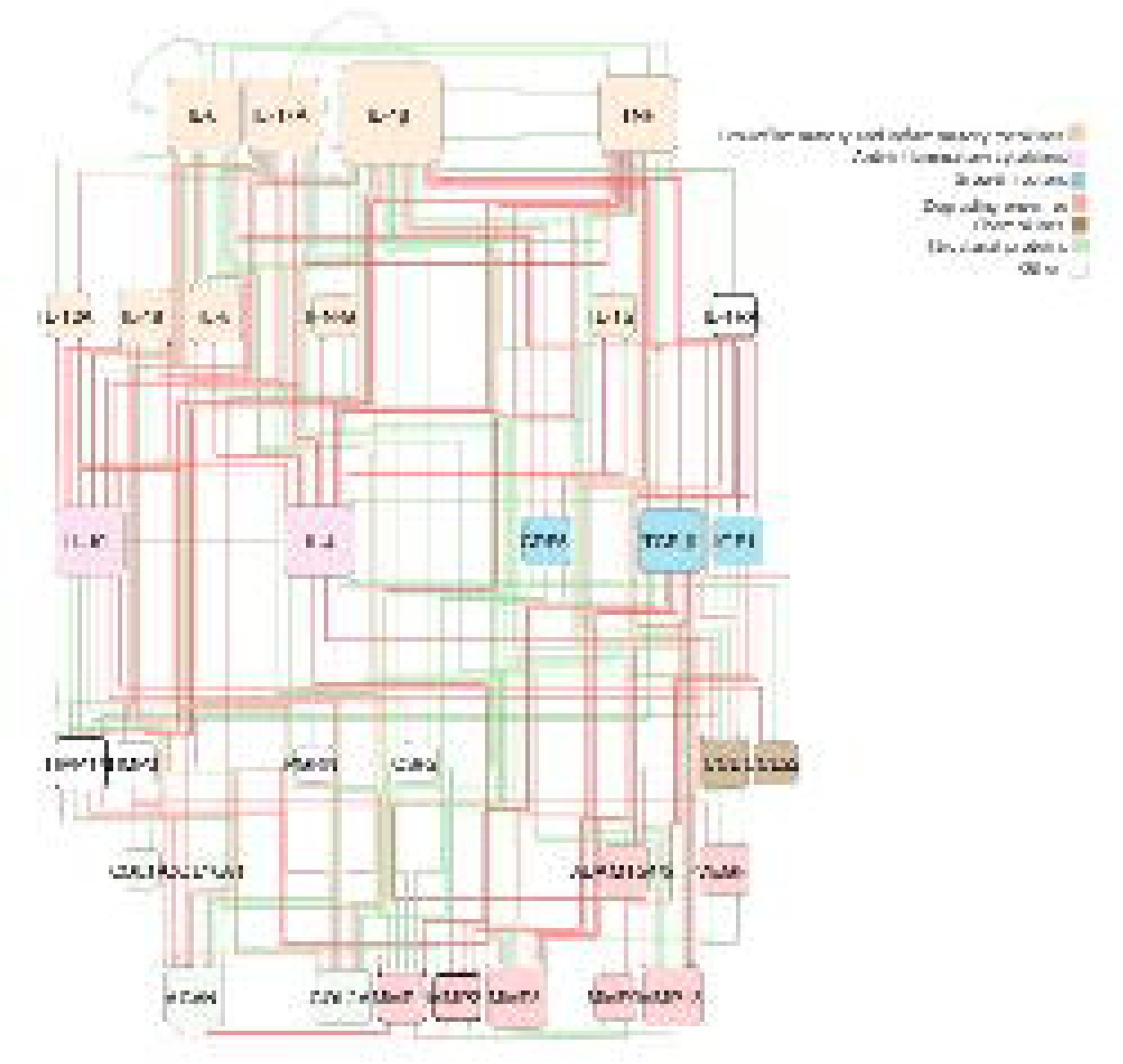
Topology representation of the enriched RNM. This network encompasses protein-protein interactions among pivotal soluble proteins involved in IVD regulation, along with general protein-protein interactions in Homo sapiens. Connections between proteins are depicted through activation (green lines) and inhibition (red lines) links. The size of each node corresponds to the number of its connections, with larger nodes indicating higher connectivity. Newly incorporated nodes are highlighted with bold borders, while bold lines denote newly added links resulting from the enrichment process.

**Figure 6.**
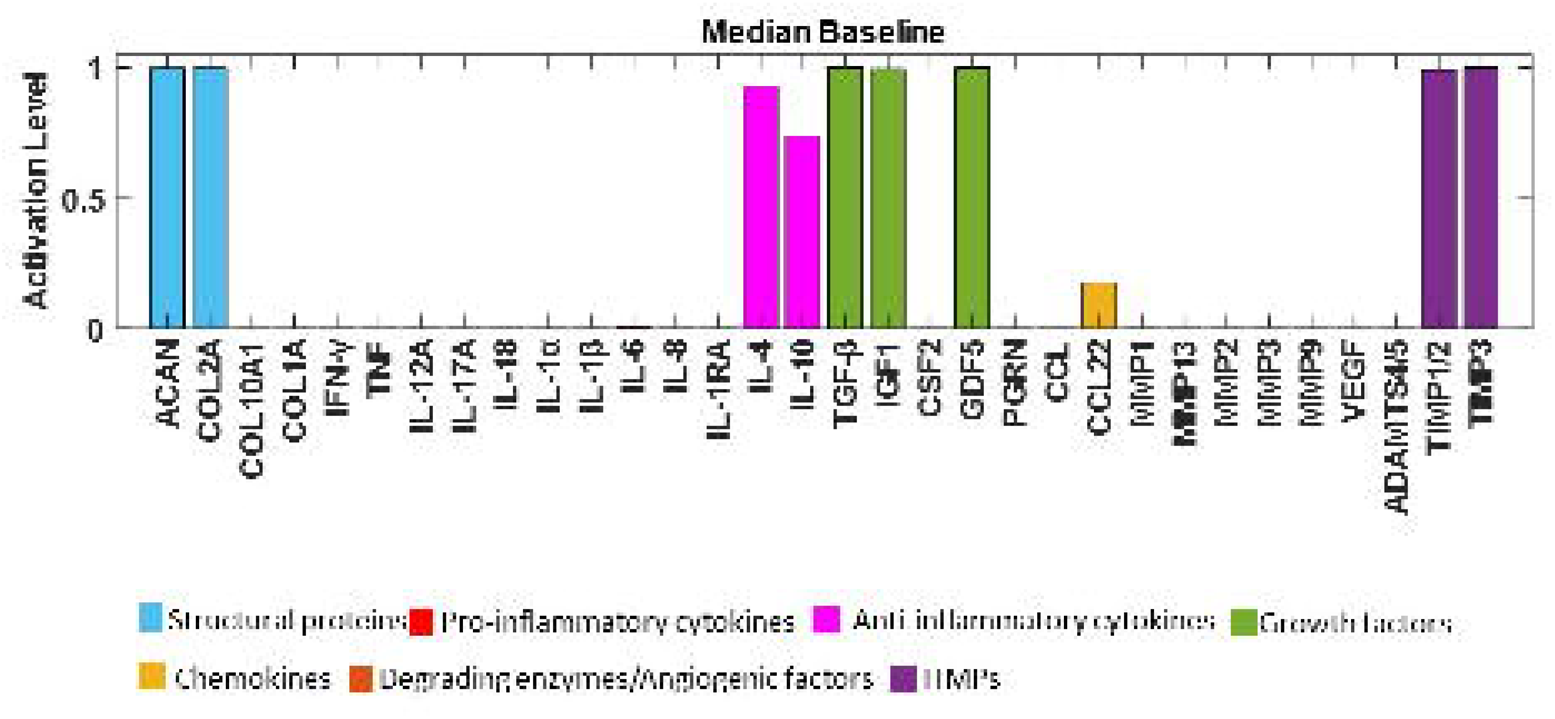
Median baseline after enrichment in the SS of the system. Activation level (0 for no activation and 1 for the highest activation) of each regulator is given in the SS of the system and are colour-labelled depending on their biochemical properties in the IVD regulation.

### Overall corpus results after enrichment

Considering the sources of knowledge out of the literature review, and after the enrichment process, 101 peer-review papers were used for the corpus. From these papers, 36 included information about outcomes of *in vitro* culture of IVD NPC, yielding a subtotal of 95 protein-protein interaction links within the network (refer to Table 1). Upon categorizing these experimental sources by the pathological state of the IVD, 54 links were derived from studies examining NPC behaviour in non-degenerate human and animal IVD tissues (as detailed in Table 2), while 50 links were informed by investigations focusing on NPC behaviour in degenerate IVD conditions.

**Table 1:**
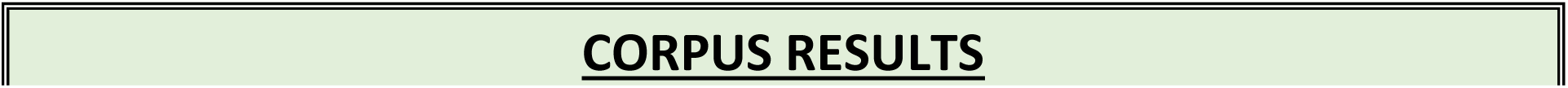
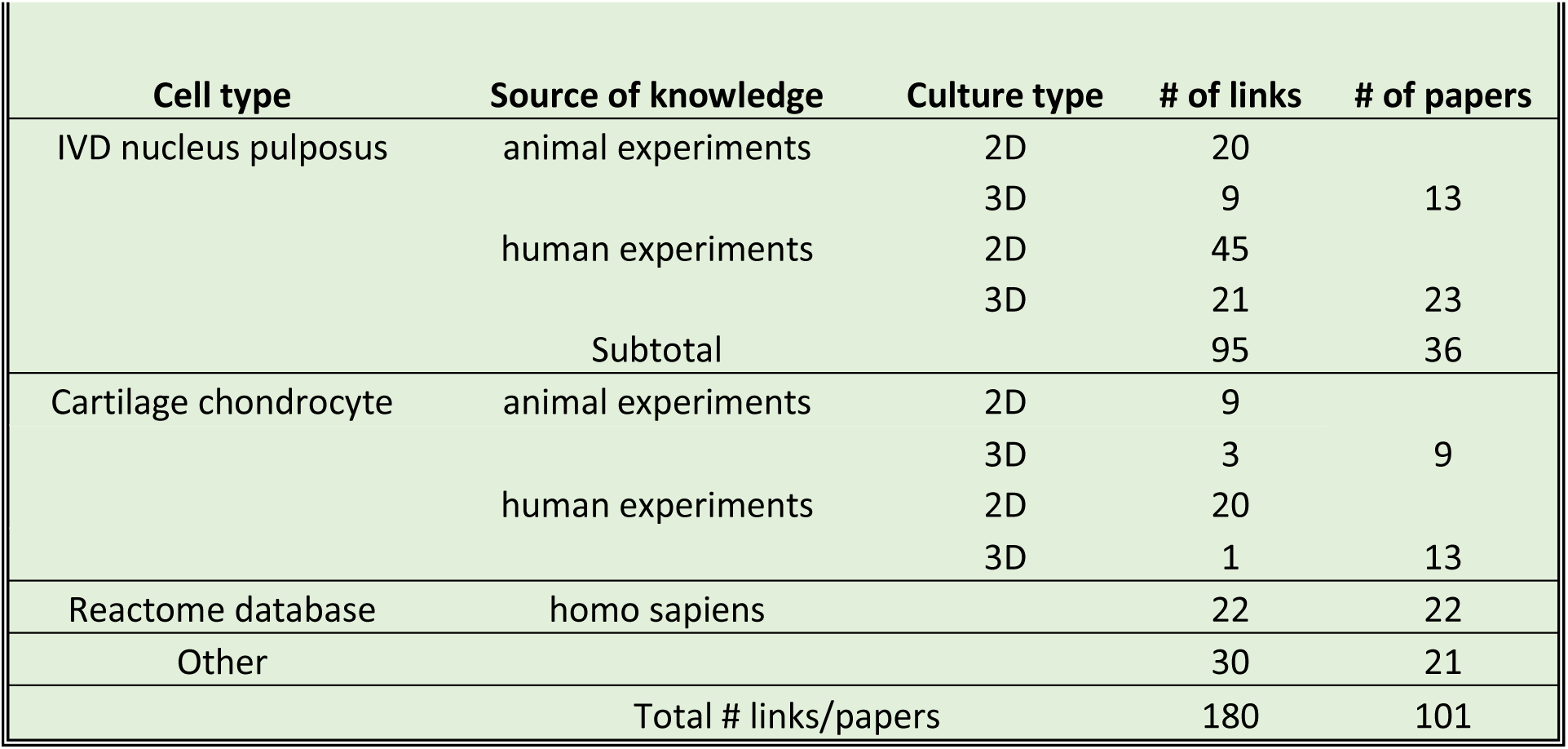
Corpus results. The cell types have been categorized as: NPC; CC; from the Reactome database enrichment; Others from manual enrichment. For each category, the source of knowledge (animal or human), the culture type (2D or 3D), the number of links (connectivity in the RNM), and the number of papers has been indicated.

**Table 2:** Pathological state results. This categorization has been made depending on the pathological state of the source material, for the NPC, and on whether cells were from humans or animals.

|  | Source of knowledge | Pathological state | # of references |
| --- | --- | --- | --- |
| IVD nucleus pulposus | human experiments | normal | 25 |
|  |  | degenerate | 50 |
|  | animal experiments | normal | 29 |

### Assessment of the network

To assess the functionality of the final system and provide an initial independent validation of the model, we tested qualitatively the enriched network against the respective results of two independent experimental treatments of human non-degenerate NPC, reported in the literature^31,32^.

Experiment 1: Network perturbation with the pro-inflammatory cytokine TNF. In accordance with findings by Tekari et al. (2022), the anticipated outcomes involved increased expressions of MMP3 and IL-6, along with decreased expressions of ACAN, COL2A, IGF1, and TGF-β. Concurrently, as demonstrated by Millward-Sadler et al. (2009), the experiments yielded a downregulation of COL2A gene expression, coupled with an upregulation of MMP3, MMP9, and MM13 gene expression. In Fig. 7A, the baseline of the enriched network is depicted by blue bars, while yellow bars represent the baseline following TNF stimulation (for boxplots see Supplementary Figure 3A). The observed behaviours align qualitatively with the anticipated outcomes from the literature. Furthermore, a Mann-Whitney U test was conducted to assess the significance of the two non-normal distributions. The analysis revealed that all changes were statistically significant (p<0.05), except for the protein PGRN.

**Figure 7.**
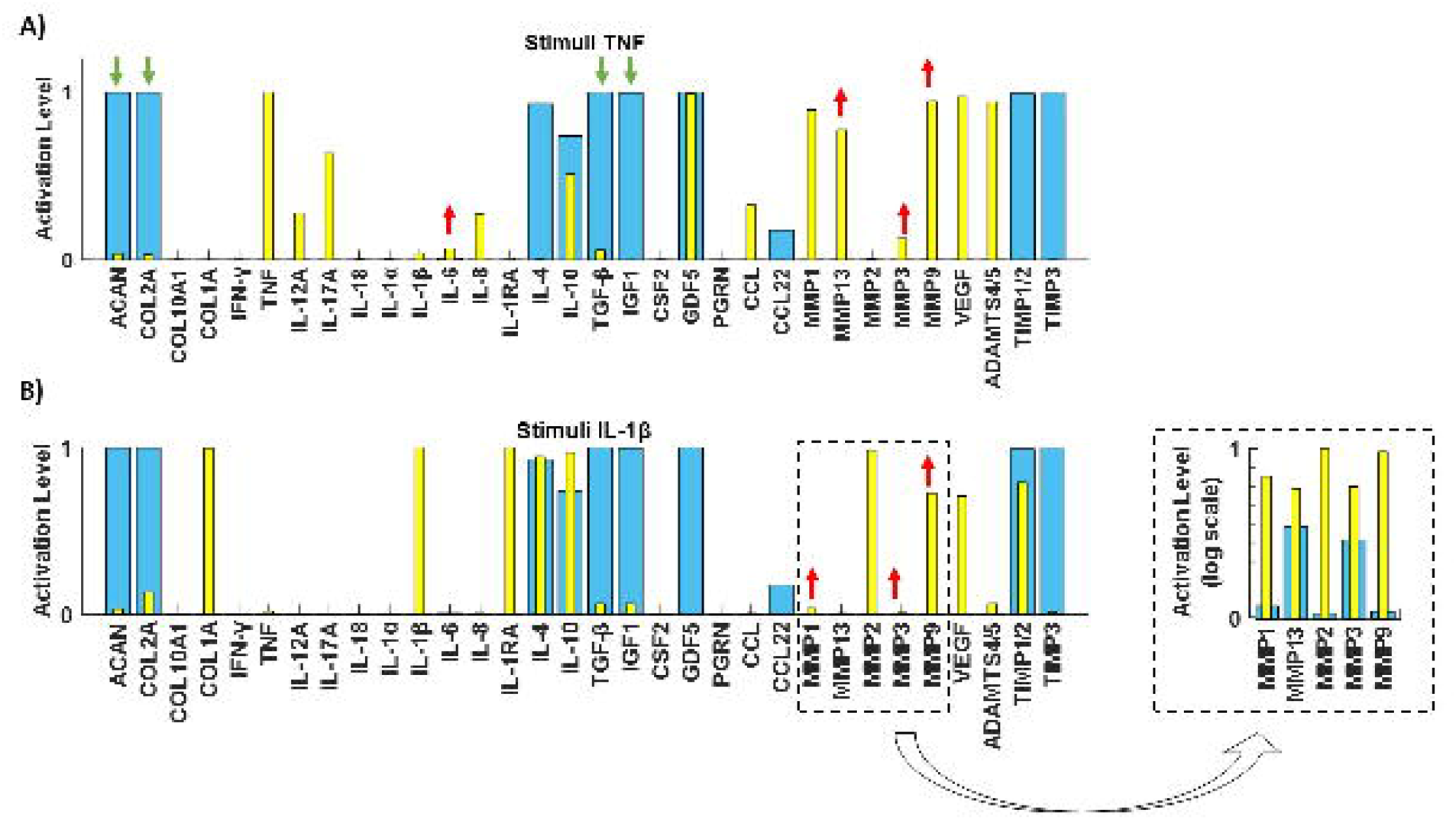
Literature-based assessment of the RNM. Testing of the RNM by replicating two independent experimental studies made in human healthy IVD NPCs. Initial SS (blue bars) and final SS after corresponding perturbations (yellow bars) with the pro-inflammatory cytokine A) TNF and B) IL-1β.

Experiment 2: Network perturbation with the pro-inflammatory cytokine IL-1β. In line with observations by Millward-Sadler et al. (2009), the anticipated responses included the upregulation of MMP3, MMP9, and MMP13 gene expression. These expected outcomes are visually represented in our system by the yellow bars in Fig. 7B (for boxplots see Supplementary Figure 3B), demonstrating the reproduction of protease responses to IL-1β stimulation. Additionally, the Mann-Whitney U statistical test confirmed the significance of all observed changes (p<0.05).

To further assess our system’s response to key anabolic regulators and explore potential rescue strategies, we simulated two distinct degenerate environments for NPC: one with pro-inflammatory cytokine TNF (Fig.8A) and another with IL-1β (Fig.8D), both set at their maximum activation levels. Under the TNF baseline, IL-4 (Fig.8B) and IL-10 (Fig.8C) significantly increased ACAN and COL2A levels (p<0.05) and decreased levels of MMPs, VEGF, ADAMTS4/5, IL-12A, IL-17A, IL-1β, IL-6, and IL-8 (p<0.05). Additionally, IL-4 significantly increased IGF1 levels (p<0.05). When using the IL-1β baseline, TGF-β significantly reduced the expression of MMPs, VEGF, and ADAMTS4/5 (p<0.05) while significantly increased ACAN and COL2A levels (p<0.05) (Fig.8E). Notably, GDF5 activation in the IL-1β baseline significantly increased ACAN and COL2A levels (p<0.05), although it did not decrease catabolic factors (Fig.8F).

**Figure 8.**
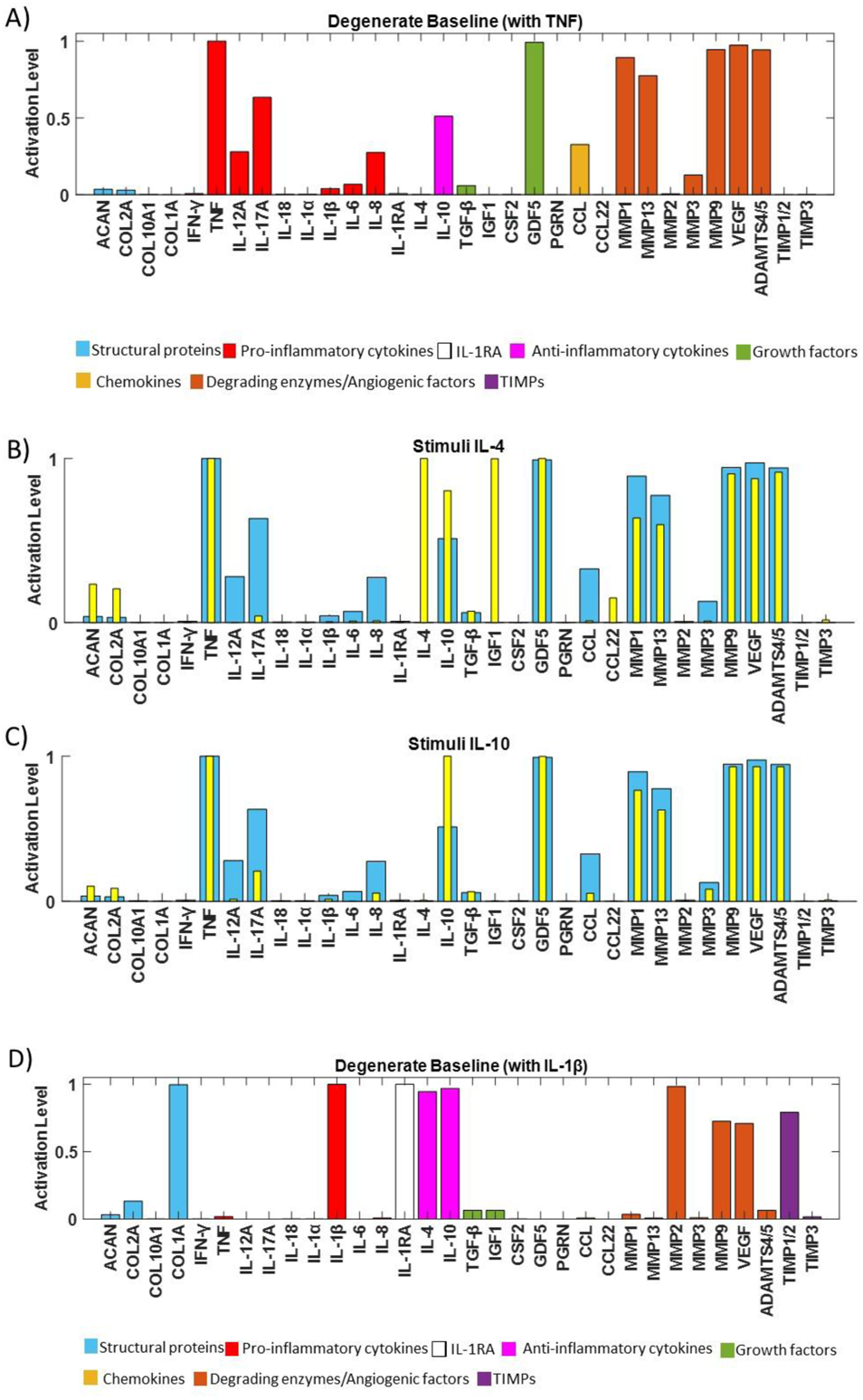
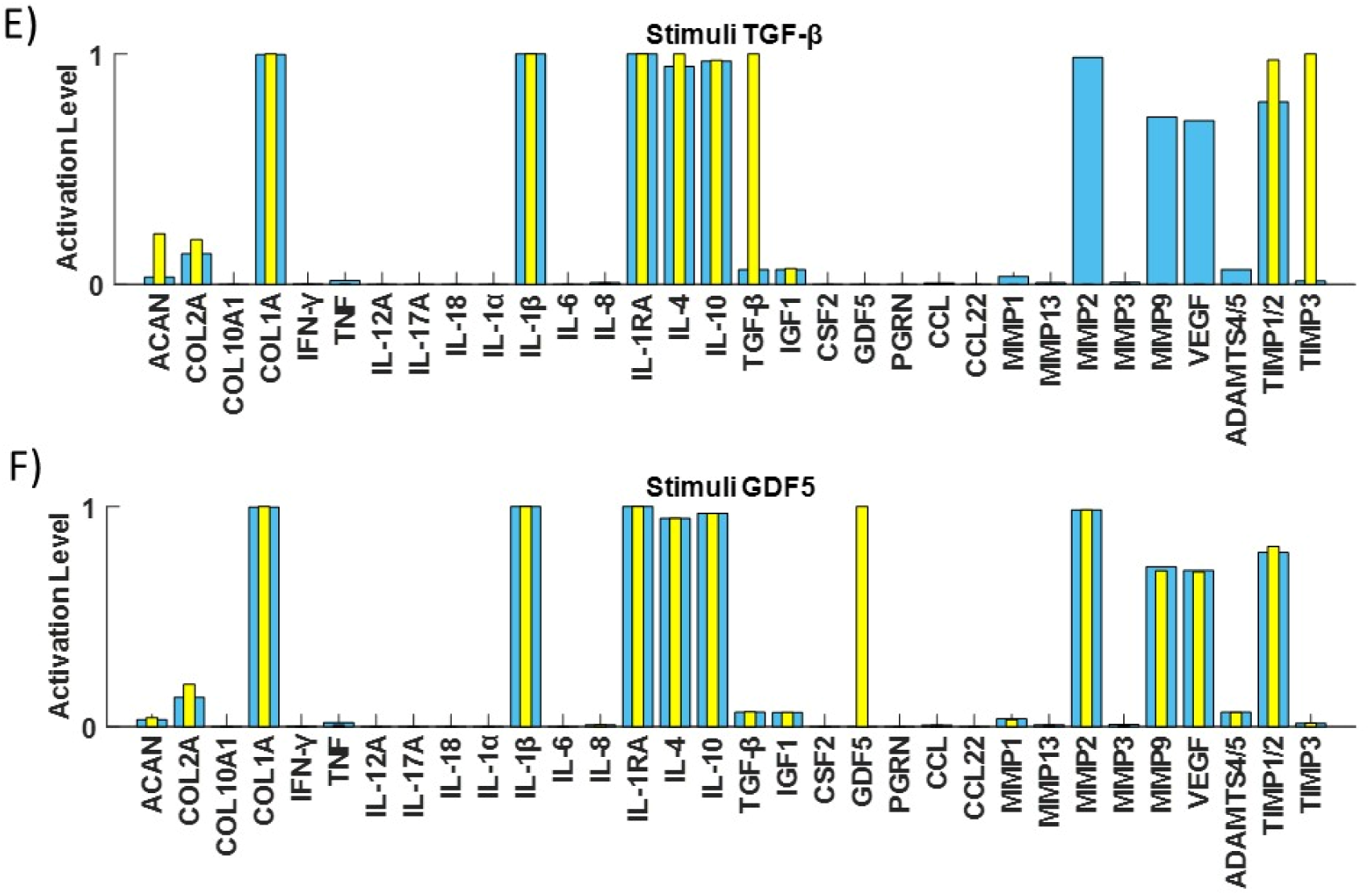
Assessment of the network through independent tests. A degenerate baseline was produced to promote catabolism by clamping TNF (Fig.8A) or IL-1β (Fig.8D) to 1. Rescue strategies were simulated by stimulating the TNF degenerate baseline with B) IL-4, C) IL-10, and the IL-1β degenerate baseline with E) TGF-β and F) GDF5 (yellow bars).

### Sensitivity Analysis

The final objective was to conduct a sensitivity analysis to identify the mediators with the most significant impact on ECM synthesis proteins (ACAN, COL1A, and COL2A) and degrading enzymes (ADAMTS4/5, MMP1, MMP2, MMP3, MMP9, and MMP13) among the 15 cytokines and 4 growth factors in the network. Simulation results revealed that among all cytokine combinations in the RNM, IL-1Ra had the highest impact on both examined groups (Fig.9A, B). Additionally, IL-6 and IL-1α showed a smaller direct effect on COL1A and COL2A, while IL-8 affected only COL2A (Fig.9A). Conversely, MMP1 and MMP2 were influenced by the inflammatory cytokines IL-6 and IL-1α (Fig.9B). Regarding the growth factors, TGF-β had the most substantial impact on the selected groups of interest (Fig.10A, B), while GDF5 significantly affected ADAMTS4/5, MMP9, and the structural proteins ACAN and COL1A (Fig.10A, B). However, VEGF and IGF1 did not exhibit any significant effect on the examined groups.

**Figure 9.**
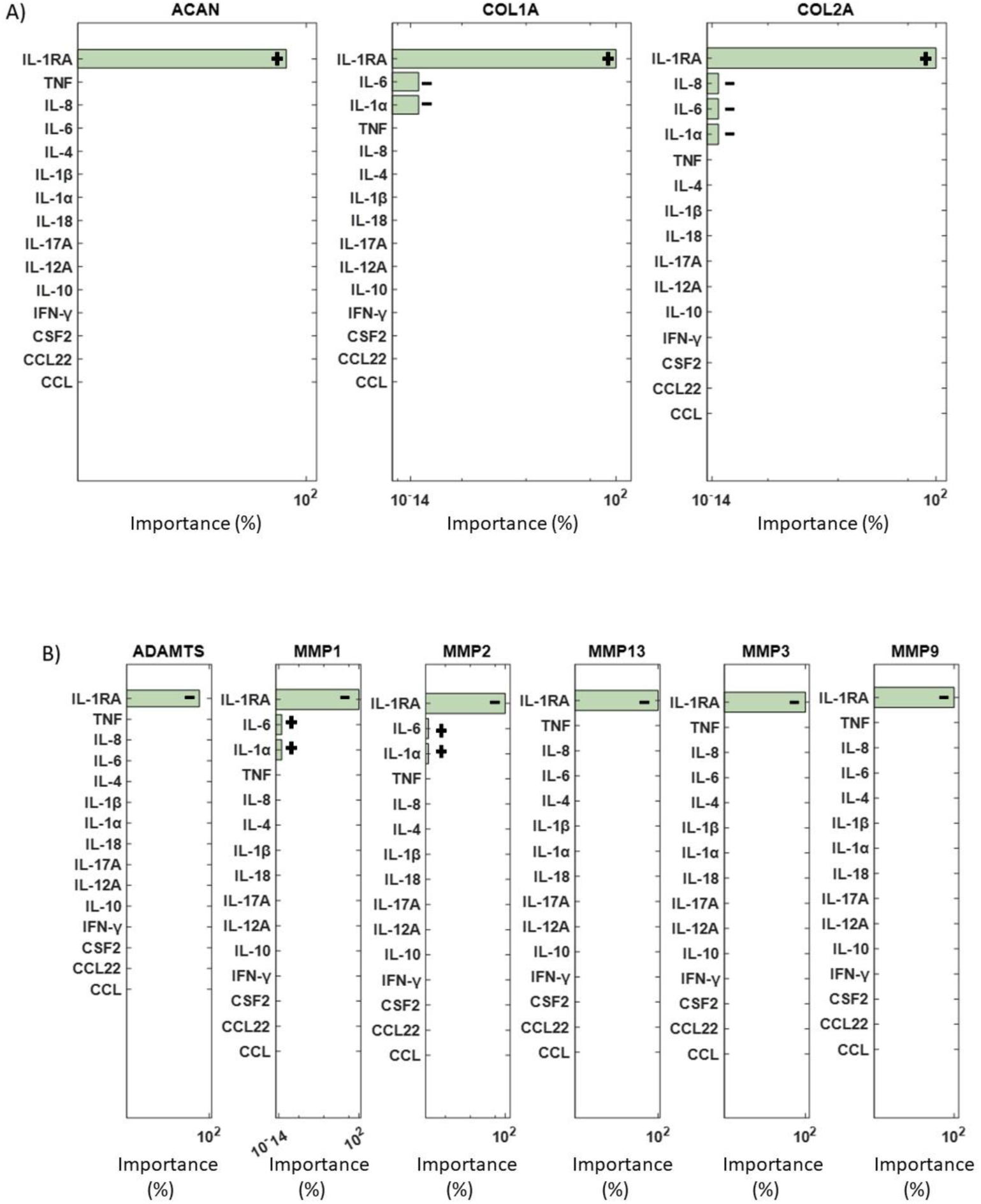
ANOVA analysis for cytokines. ANOVA analysis results showing the most significant cytokines (green bars) that affected A) the structural proteins and B) the degrading enzymes are represented by green bars. Non-significant combinations are indicated by pink bars, while combinations that fall outside the range of significance are not depicted. +: positive impact, -: negative impact.

**Figure 10.**
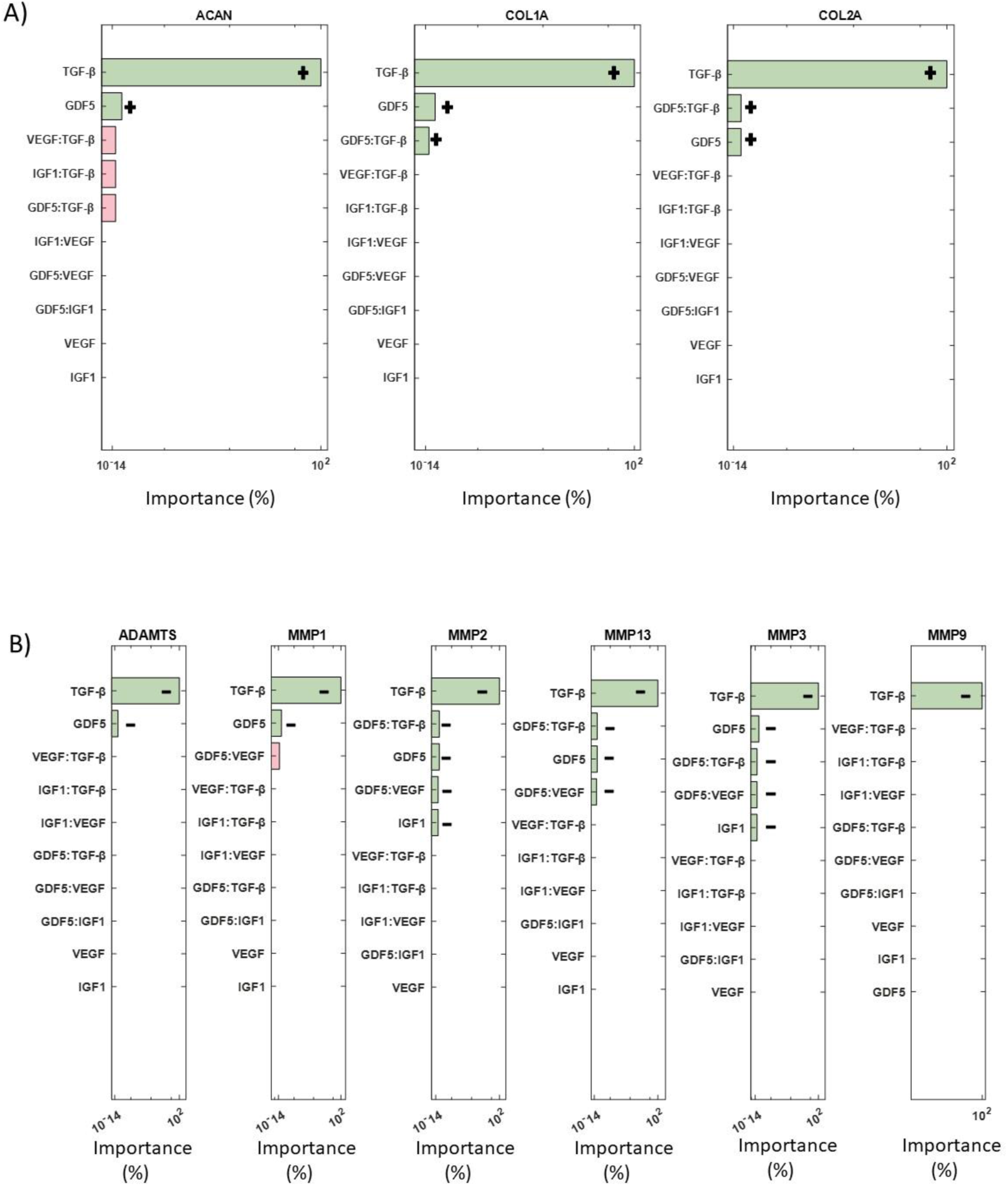
ANOVA analysis for growth factors. Significant combinations of growth factors that impact A) structural proteins and B) degrading enzymes are represented by green bars. Non-significant combinations are indicated by pink bars, while combinations that fall outside the range of significance are not depicted. +: positive impact, -: negative impact.

Fig.9 ANOVA analysis for cytokines. ANOVA analysis results showing the most significant cytokines (green bars) that affected A) the structural proteins and B) the degrading enzymes are represented by green bars. Non-significant combinations are indicated by pink bars, while combinations that fall outside the range of significance are not depicted. +:positive impact, -:negative impact.

## Discussion

The disruption of a balanced regulation of the ECM by IVD cells might cause or accelerate the degeneration of the disc. Biochemical factors need to be explored carefully, not only to understand the mechanisms that contribute to ECM degradation, but also to predict and/or prevent the triggering of these mechanisms. Multiple experimental measurements have generated valuable knowledge about IVD protein response against pro-catabolic, anti-catabolic and growth factor stimulations, but the integration of such knowledge is still missing, as targeting only a single protein or few proteins at once is not sufficient to test the effect of the many proteins that can shape the biochemical microenvironment of NPC. Network modelling can provide a holistic and intuitive vision of the complex behaviour and interactions among proteins, and complement *in-vitro* studies, as it allows to explore perturbations/interactions that would require further experimental exploration.

Different approaches can be adopted to build a network model: either gather literature information about the biochemical IVD regulation (knowledge-driven network) or exploit directly experimental measurements (data-driven network), as performed by Baumgartner et al. (2021)^26^. While the latter can predict IVD cell responses in a pro-inflammatory environment, in terms of protein expression, it is limited on TNF and IL-1β stimulations and it did not consider important pro-anabolic and pro-catabolic factors in IVD research. On the other hand, our model provides a unique corpus of knowledge able to represent a relatively large spectrum of NP cell activity predictions upon different perturbations. In addition, it can test virtually perturbations still not experimentally explored, which facilitates, and identifies priorities for future experimental study designs.

The initial system exhibited an anticipated catabolic response, given that 52% of the experimental samples sourced from the literature either originated from degenerated discs or were artificially induced into a degenerative state prior to testing. However, our primary aim was to construct a network that accurately represented a non-degenerate IVD, enabling us to evaluate how various stimuli influence metabolic shifts. Consequently, enriching the network with relevant protein-protein interactions, particularly those aligned with our objectives, was deemed a logical progression. By incorporating key missing interactions (depicted as bold lines in Fig. 5), we successfully attained an anabolic state. This state is distinguished by heightened activation levels of essential ECM components such as ACAN and COL2A, along with pivotal growth factors including GDF5, IGF1, and TGF-β. Conversely, activation levels of angiogenic factors (e.g., VEGF), catabolic markers (MMPs, ADAMTS4/5), and pro-inflammatory cytokines (such as interleukins, TNF, and IFN-γ) were notably suppressed. This iterative process underscored not only the efficacy of our approach but also the imperative for further experimental studies specifically focused on IVD biology. Out of the 101 papers analysed, 36 pertained to IVD NP cells, 22 focused on cartilage chondrocyte cells, while the remaining 43 addressed other cell types, highlighting the pressing need for more comprehensive investigations within the field.

The significance of enrichment and subsequent network predictions finds robust support in the experiments conducted by Li et al. (2014)^33^. The authors investigated the impact of treating degenerate NP canine cells with IL-10 and TGF-β, revealing a notable suppression in the release of inflammatory cytokines IL-1β and TNF^33^. Such effect shall further limit the additional catabolic expression of proteases and deterioration of the ECM^34,35^. Ge et al. (2020) showed that IL-10 treatment on NP degenerate cells could significantly increase the protein expression of type II collagen and aggrecan while decreasing protein levels of type X collagen and endogenous mRNA expression of TNF and IL-1β^12^. A study that shows the importance of our enriched regulators (IL-4, TIMP1/2, and TIMP3), was demonstrated by Kedong et al. (2020), who determined the anabolic role of the anti-inflammatory IL-4 against the inflammatory markers IL-6 and IL-8^11^ while TIMPs act as a balance factor to the catabolic behaviour of MMPs and ADAMTs^36,37^. Notably, TIMP3 emerges as a potent inhibitor of ADAMTS4 and ADAMTS5, playing a pivotal role in safeguarding against ECM matrix-degrading enzymes^38^. Moreover, despite the concurrent increase of TIMP1, a major MMP inhibitor, alongside MMPs during disc degeneration, TIMP3 exhibits a distinct behaviour, not following the trajectory of cell immunopositivity for ADAMTS during degeneration^39^. Lastly, the positive outcomes of enrichment are further exemplified by the addition of IL-1Ra, the natural inhibitor of IL-1^40^. When delivered through gene therapy, this receptor antagonist could significantly diminish the expression of MMP1, MMP3, MMP13 and ADAMTS4, as well as the activity of these enzymes in the NP and the AF, by more than 95%^40^.

In our pursuit of independently assessing the enriched network, we conducted a thorough literature search to identify experimental studies not initially incorporated into the knowledge representation that formed the network topology. The literature was filtered to solely target stimulated NP cells derived from non-degenerate human IVD samples and build a set of tests as relevant to human IVD biology, and coherent, as possible. First, we stimulated the network with the pro-inflammatory cytokine TNF. This stimulation prompted a significant catabolic response within our system, evidenced by the upregulation of IL-6 (p < 0.05) and MMP3 (p < 0.05), accompanied by the downregulation of ACAN (p < 0.05), COL2A (p < 0.05), IGF1 (p < 0.05), and TGF-β (p < 0.05). Notably, this trend aligned with findings reported by Tekari et al. (2022) as well^31^. When comparing the nodal activation level under TNF stimulation with the second experimental study from Millward-Sadler et al. (2009)^32^, we observed consistent upregulation of MMP3 (p < 0.05), MMP13 (p < 0.05), IL-1β (p < 0.05), and IL-1Ra (p < 0.05), alongside downregulation of COL2A (p < 0.05) and COL1A (p < 0.05). Secondly, we perturbated the network with the pro-inflammatory cytokine IL-1β, to confirm the upregulation of MMP3 (p<0.05), MMP13 (p<0.05), MMP9 (p<0.05), and TNF (p<0.05) as measured from Millward-Sadler et al. (2009)^32^. Essentially, our *in-silico* simulations corroborated these experimental results, providing further validation of our enriched networks’ predictive capabilities.

In another aspect of our assessment, we sought to explore the potential effects of IL-4, IL-10, TGF-β, and GDF5, all recognized for their reported anabolic responses in a degenerative environment^11,12,33,41–44^. To establish a degenerate baseline, we used two approaches: clamping TNF at a level of 1 and clamping IL-1β at a level of 1. These cytokines are proposed to play a key role in intervertebral disc degeneration (IDD), as indicated in previous studies^4,32,45,46^. Baseline characterized by elevated expression levels of MMPs, ADAMTS4/5, and VEGF, all associated with extracellular matrix (ECM) degradation and angiogenesis^32,39,47,48^, were thus created. We then tested the effects of anti-inflammatory cytokines and growth factors on these baselines. We saw that anti-inflammatory cytokines could rescue the IDD when applied in an environment like the TNF baseline, while growth factors (TGF-β better than GDF5) would promote anabolism when applied in environment similar to IL-1β baseline.

More specifically, Ge et al. (2020) has previously demonstrated that IL-10 could be used to delay IDD through its anti-inflammatory response by inhibiting p38 MAPK pathway activation^12^, while recently Kedong et al. (2020) has proposed that the recombinant IL-4 delivery into the IVD might have a beneficial therapeutic effect by reducing disc inflammation following herniation^11^. In our *in silico* simulations, IL-4 decreased MMP3 and IL-17A, consistent with *in vitro* results from Chambers et al. (2013)^49^ and Sandoghchian et al. (2011) ^50^. Furthermore, IL-4 increased ACAN, COL2A, and IGF1, aligning with experimental results in chondrocyte cells^51,52^. IL-10 also increased COL2A, a result observed experimentally in NPC^12^ and decreased pro-inflammatory cytokines (IL-6, IL-8, IL-17A, IL-12A), corroborating findings in other cell types^53,54^. However limited studies have investigated the effects of IL-4 and IL-10 on human NP cells from non-herniated material which could be infiltrated with immune cells. TGF-β stimulation led to depletion of MMP2 expression, a result reported previously in bovine IVD NP cells by Pattison and colleagues^55^. It also eliminated the catabolic markers MMP3, MMP9, MMP13, and ADAMTS4/5, whose activity is closely related to ECM degradation^5^. Although the downregulation of these markers by TGF-β has been observed in general human cell types^27,56^, such findings are lacking in NP cells, underscoring the need for further experimental studies specific to IVDs. Notably, TGF-β induced a high expression level of TIMP3, a result not previously reported in NP cells but observed in fibroblasts and chondrocytes^57,58^, which share similar functional and cellular characteristics with NP cells^59–61^. Additionally, TGF-β significantly increased structural proteins ACAN and COL2A, consistent with findings in NPCs by Masuda et al. (2003)^62^. GDF5, while not significantly altering the degenerate state, did decrease TNF expression, consistent with findings by Guo et al. (2021)^42,63^, suggesting GDF5’s efficacy in suppressing ECM degradation. Our *in silico* model also promoted anabolism by increasing ACAN and COL2A, as found by Chujo et al.^14^ and Maitre et al. (2009)^43^.

Here, we also investigated the potential rescue effects of IL-4, IL-10, TGF-β, and GDF5 in a degenerate baseline generated from IL-1β and TNF co-stimulation. Only TGF-β was able to decrease the catabolic effects on the MMPs, while no significant changes were observed in structural proteins to promote anabolism (Supplementary Figure 4). The expression of TNF and IL-1β in degenerate IVDs is complex and can vary independently. It is unlikely that IVDs would express only one cytokine (IL-1β/TNF), however depending on the stage of degeneration and infiltration of inflammatory cells following AF and CEP fissures, it is possible that some discs could have differential levels ^32,64^. Understanding these dynamics is crucial for developing targeted therapies for disc degeneration.

The sensitivity analysis aimed to determine key determinants of the ECM regulation in the disc. Among the cytokines the anti-catabolic factor IL-1Ra had the most impact, showing the importance of the natural IL-1 inhibitor, whose deficiency might develop IDD ^65^. During disc degeneration COL2A and ACAN are decrease^66^, which is associated with dehydration of the disc and reduced ability to withstand loads. Thus, if IL-1Ra via the inhibition of native IL-1 agonists (α and β) could prevent the decrease in matrix synthesis, which accompanied with growth factor stimulation, either native or applied could promote new matrix synthesis, IDD could be halted, and potential regeneration induced. However, the realization of such therapeutic potential hinges upon a deeper understanding of IL-1Ra’s regenerative properties, necessitating further experimental investigations. Moreover, sensitivity analysis underscored the profound impact of transforming growth factor-beta (TGF-β) on both structural proteins and matrix-degrading enzymes. This finding emphasizes the pivotal role of TGF-β in orchestrating ECM homeostasis and suggests its potential as a therapeutic target for IDD management. Moreover, *in-silico* co-stimulation of TGF-β with IL-1β, and TNF was able to reduce the presence of pro-inflammatory cytokines and increase ACAN, explaining its strong impact. These insights not only deepen our understanding of the molecular mechanisms underlying disc degeneration but also highlight promising avenues for therapeutic intervention aimed at promoting disc regeneration.

To conclude, the proof of concept development of an *in silico* network model demonstrates the importance of a baseline in network modelling. We have achieved a unique corpus of IVD biochemical regulators and a baseline which is proposed to represent the non-degenerate state of the disc. Our network showed that TGF-β is promising in the regeneration of the disc, while more exploration of the IL-4 and IL-10 anti-inflammatory effects is needed. Considering rescue strategy simulations, the combination of autologous anti-inflammatory serum containing TGF-β, GDF5, IL-10, and IL-4 emerges as compelling potential biological treatment for IDD. In addition, the initial state of the nodes plays a very important role in the response of the system against stimulations and for that more IVD specific experimental studies should be conducted. Current treatments for IDD are mostly conservative and can only alleviate the pain, but not regenerate the disc^67^. It is essential to acknowledge that while our model provides valuable insights, it does not capture intracellular changes or include pathways, which are crucial for comprehensive analysis. Future iterations of our regulatory network model should incorporate these elements to serve as a versatile tool for guiding experimental studies, predicting outcomes, and testing IDD drugs.

## Methods

### Corpus

As a first step, we searched in the open database PubMed for IDD chondrocyte protein-protein interactions. Specifically, our search terms included the following combinations:

`o` (Intervertebral disc degeneration) AND (nucleus pulposus cell)
`o` (Intervertebral disc degeneration) AND (cytokine)
`o` (Intervertebral disc degeneration) AND (anti-inflammatory cytokine)
`o` (Intervertebral disc degeneration) AND (growth factor)
`o` (Intervertebral disc degeneration) AND ((Matrix metalloproteinases) OR(MMP))
`o` (Intervertebral disc degeneration) AND ((a disintegrin and metalloproteinase with thrombospondin motifs) OR(ADAMTS))
`o` (Intervertebral disc degeneration) AND (chemokine)

From the plethora of articles identified, a meticulous review yielded 34 peer-reviewed studies that formed the foundation for constructing our initial IVD regulatory network model. This compilation encompassed data pertaining to direct activation, inhibition, upregulation, and downregulation effects among a diverse array of molecular entities crucial in regulating IVD function. These entities spanned structural proteins, degrading enzymes, angiogenic factors, pro-inflammatory, inflammatory, and anti-inflammatory cytokines, as well as chemokines. In refining our dataset, we opted to streamline the representation of C-C Motif Chemokine ligands by merging them into a single category denoted as CCL. However, we noted a distinct behaviour exhibited by CCL22 compared to others within this category. While IL-1Β activated other CCLs, it paradoxically inhibited CCL22^68^, warranting its separation from the merged group. This final dataset is accessible in the corresponding repository^28^. Leveraging the Cytoscape_v3.8.2 platform, we employed a node-edge visualization approach to depict proteins as nodes and illustrate their activation and/or inhibition effects through connecting edges. This strategy offers an intuitive and comprehensive representation of the intricate regulatory mechanisms governing IVD biology, facilitating in-depth analysis and interpretation of IDD-related protein interactions.

### Semi-quantitative system resolution

To transition the knowledge-based regulatory network (RN) from a static snapshot to a dynamic representation, we employed a method proposed by Mendoza, which utilizes fuzzy logic semi-quantitative interpolation of boolean solutions^20^. This approach enables the conversion of boolean logic into ordinary differential equations, facilitating the modelling of dynamic behaviour within the network.

The calculation of the final expression for each node involves a set of ordinary differential equations, with the dynamics described by the following equation:

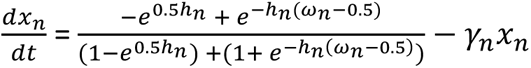

Here, the left-hand side represents the activation function, while the right-hand side depicts a decay function proportional to the node x_n_, weighted by a factor γ>0. The activation term incorporates the sigmoid function ω, which computes the total input of a node x_n_.

Generally, each node has different connectivities, according to the number of activators and/or inhibitors (Fig.11). Thus, ω is given by the following equations, each of them describing the possibility of each node having solely activators, solely inhibitors or a combination of both. The equation is as follows:

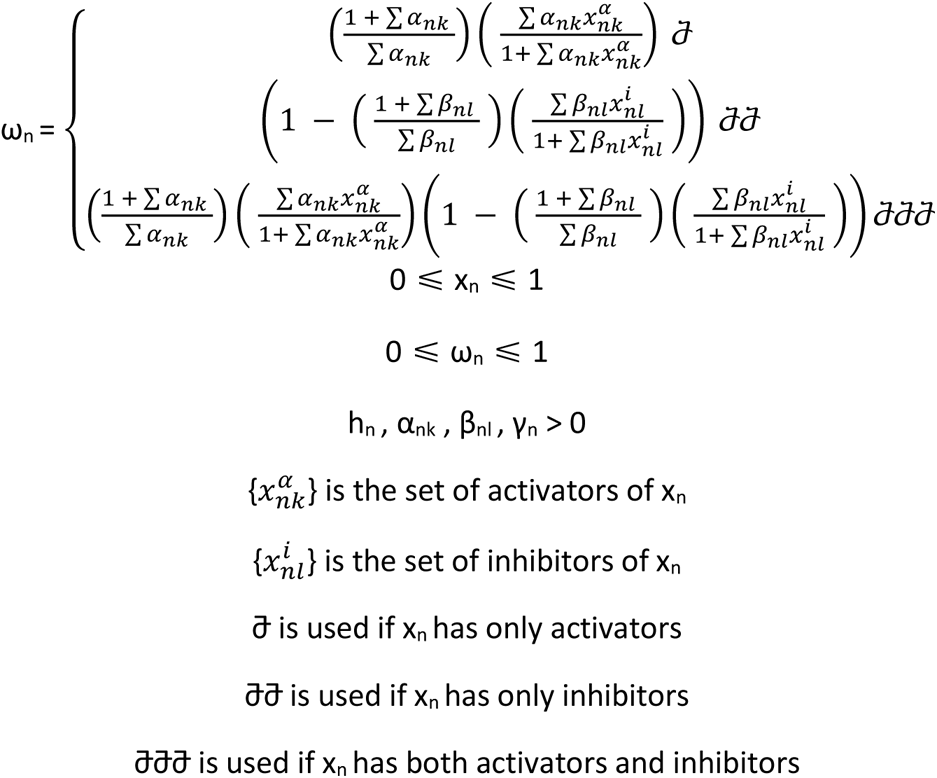

**Figure 11.**
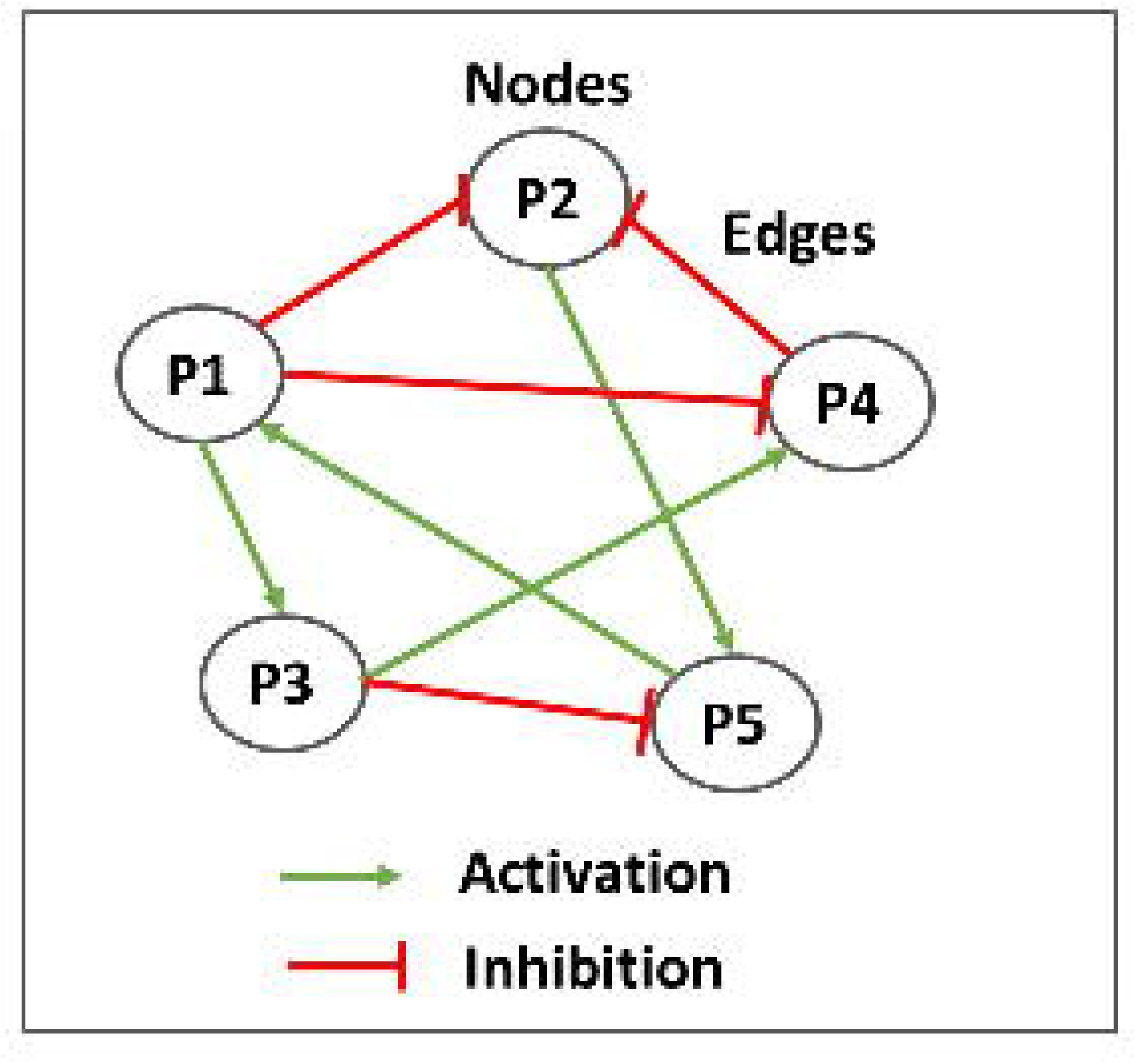
Simplified representation of a regulatory network. In a regulatory network model, proteins are represented by nodes that interact among each other through activation (red lines) and/or inhibition (green lines) edges.

In these equations α>0 and β>0 are the weights of activators and inhibitors, respectively and take the values of α=1, β=1 for simplicity as in spite of their values, they do not affect the monotonically behaviour of the ω function between its intervals. Finally, we choose h=10 and γ=1 for all nodes, as proposed by Mendoza et al. (2006), a choice serving the convenient and smooth biological behaviour of the system.

The system was resolved using the fourth-order Runge-Kutta method in Matlab 2018b (The MathWorks, Inc, Massachusetts, USA) with the ode45 solver to determine the global attractor or steady state (SS) of the system (Fig.12). This SS represents a practical approximation of an artificial time period within a cellular process, likely confined to a localized region of interest. Consequently, we designated this SS as the baseline of each protein within the network. To ensure the robustness of our baseline calculations, we conducted 100 simulations for each protein, varying initial conditions based on Mersenne Twister 0 seeds. This rigorous approach allowed us to confirm the consistency of the baseline across various initial conditions. Additionally, it ensured that each node converged to a steady state, reflecting the system’s stable behaviour. Upon examination, it became evident that the steady state (SS) values for each protein did not conform to a normal distribution pattern; rather, the distribution appeared skewed. In response to this observation, we opted to employ a boxplot visualization technique to illustrate all SS values comprehensively. This approach offers a detailed representation of the baseline, capturing the distribution of SS values along with key statistical metrics such as median, quartiles, and outliers. However, owing to the limited variance in the values and the skewed distribution, boxplots were not sufficiently sensitive in highlighting significant differences. To enhance clarity and facilitate better interpretation for readers, we chose to present the median of the baseline using barplots. Meanwhile, the detailed boxplot analyses, encompassing additional statistical insights, can be accessed in the Supplementary Figure 1 and Supplementary Figure 2. This approach ensures that readers can easily discern the baseline trends while still having access to comprehensive complete datasets provided by the boxplots. The code can be found in the following repository^29^.

**Figure 12.**
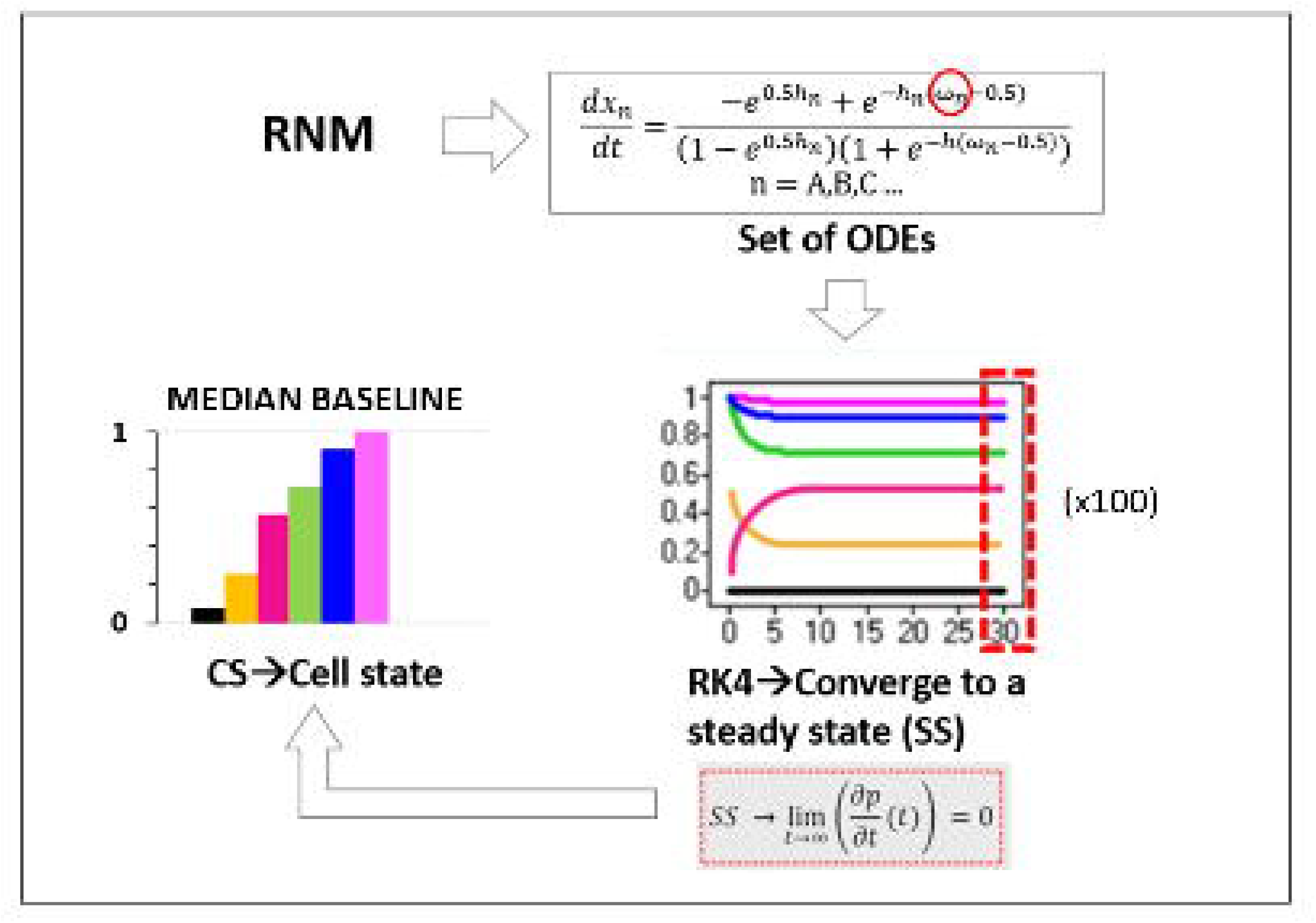
Schematic representation of the semi-quantitative method employed to analyse the regulatory network model (RNM). The dynamics of the RNM were captured through a system of ordinary differential equations (ODEs), where the activation of each node is determined by the number of inhibitors and activators. The ODEs were solved using the fourth-order Runge-Kutta method (RK4), initialized with random system conditions. After 30 seconds, the system converged to a steady state (SS), serving as the baseline for each protein within the network. Notably, 100 simulations were conducted, and all resulting SS values were utilized for subsequent analysis, as illustrated by the boxplots.

### Enrichment

The enrichment of the regulatory network was done firstly through the STRING (Version 11.5) public database. This procedure retrieved information about protein-protein interactions which we could not find in literature when looking specifically for IVD NPC regulation. Text mining, experiments and databases were selected as interaction sources while the interaction score with highest confidence of 0.900 was applied in the direction of having strong and reliable biological information. We examined extensively one by one all the links among 34 proteins that already existed in the initial RNM to enrich it with the missing ones. Additionally, a manual enrichment process was conducted by scouring PubMed for information on growth factors such as TGF-β and IGF, as there were still gaps in the network regarding these factors. Furthermore, to ensure comprehensive coverage of key proteins implicated in IVD pathogenesis, additional proteins were manually added including MMP1, MMP2, TIMP1, and IL-1Ra based on recent investigations documented in the PubMed literature. These proteins play pivotal roles in IVD pathophysiology, and their inclusion was deemed essential for a thorough understanding of regulatory network dynamics.

For the manual enrichments the keywords in PubMed were:

`o` ((intervertebral disc degeneration)) AND ((transforming growth factor beta) OR (TGF-β)) (20/10/2021)
`o` ((intervertebral disc degeneration)) AND ((insulin-like growth factor) OR (IGF)) (20/10/2021)
`o` ((intervertebral disc degeneration)) AND ((Matrix Metallopeptidase 1) OR (MMP1)) (20/10/2021)
`o` ((intervertebral disc degeneration)) AND ((Matrix Metallopeptidase 2) OR (MMP2)) (20/10/2021)
`o` ((intervertebral disc degeneration)) AND ((tissue inhibitor of metalloproteinases) OR (TIMP1)) (20/10/2021)
`o` ((intervertebral disc degeneration)) AND ((Interleukin-1 receptor antagonist) OR (IL-1Ra)) (20/10/2021)

As a last step of the enrichment, we used missing links found in osteoarthritis chondrocytes regarding TIMP1, TIMP2, TGF-β, IL-10 and IL-4 ^27^, as osteoarthritis is a disease with common mechanisms as IDD this was a similar pathway and cell type to enhance the network. Detailed information for the enriched network used can be found in the corresponding repository^28^. In the final enriched network ADAMTS4 and ADAMTS5 were merged as ADAMTS4/5, TIMP1 and TIMP2 as TIMP1/2, while IL-23A and IL-32 were removed to simplify our network as they showed limited connections. Eventually, the stable SS of the enriched network was solved by using the method explained in the semi-quantitative system resolution (Fig.12) as done for the knowledge-based network initially.

### Assessment of the network

After building the network we evaluated its performance by exploring the anabolic or catabolic effect of each protein in IVD regulation (database). A widely accepted principle dictates that a high activation level of structural proteins such as COL2A and ACAN, as well as growth factors, coupled with a low activation level of proteases, ADAMTS, and pro-inflammatory cytokines, promotes anabolism within the IVD. Conversely, a low activation level of structural proteins and growth factors, along with a high activation level of MMPs, ADAMTS, and pro-inflammatory cytokines, indicates a catabolic state.

The experimental system was further tested by comparison to independent experiments from the literature, by replicating theoretically two in-vitro experiments. More specifically, in the first test according to Tekari et al. (2022) ^31^, in which they used non-degenerate human IVD NP cells collected from trauma patients undergoing spinal fusion surgery without any history of disc degeneration before the operation, the current *in silico* network was stimulated with the pro-inflammatory cytokine TNF. In the second test, according to Millward-Sadler et al. (2009)^32^, in which they used non-degenerate and degenerate human NP cells obtained from post-mortem samples or from surgery, and graded histologically according to the method of Sive and colleagues^69^, we stimulated the network with the pro-inflammatory cytokines TNF and IL-1β. In both tests, we analysed the behaviour of our network post-stimulation and compared it with the results obtained from the corresponding experiments. This comparison allowed us to evaluate the functionality and accuracy of our network model in replicating experimental biological responses.

### Sensitivity Analysis

A sensitivity analysis was conducted to identify the mediators that most significantly affected the system (Fig. 13). Specifically, the analysis aimed to determine which cytokines/chemokines (CSF2, IFN-γ, IL-1α, IL-1β, Il-1Ra, IL-4, IL-6, IL-8, IL-10, IL-12α, IL-17A, IL-18, TNF, CCL, CCL22,), growth factors (GDF5, IGF1, TGF-β, VEGF), and their second-order interactions significantly impact the groups of structural proteins (ACAN, COL1A, COL2A) and degrading enzymes (ADAMTS4/5, MMP1, MMP2, MMP3, MMP9, MMP13).

**Figure 13.**
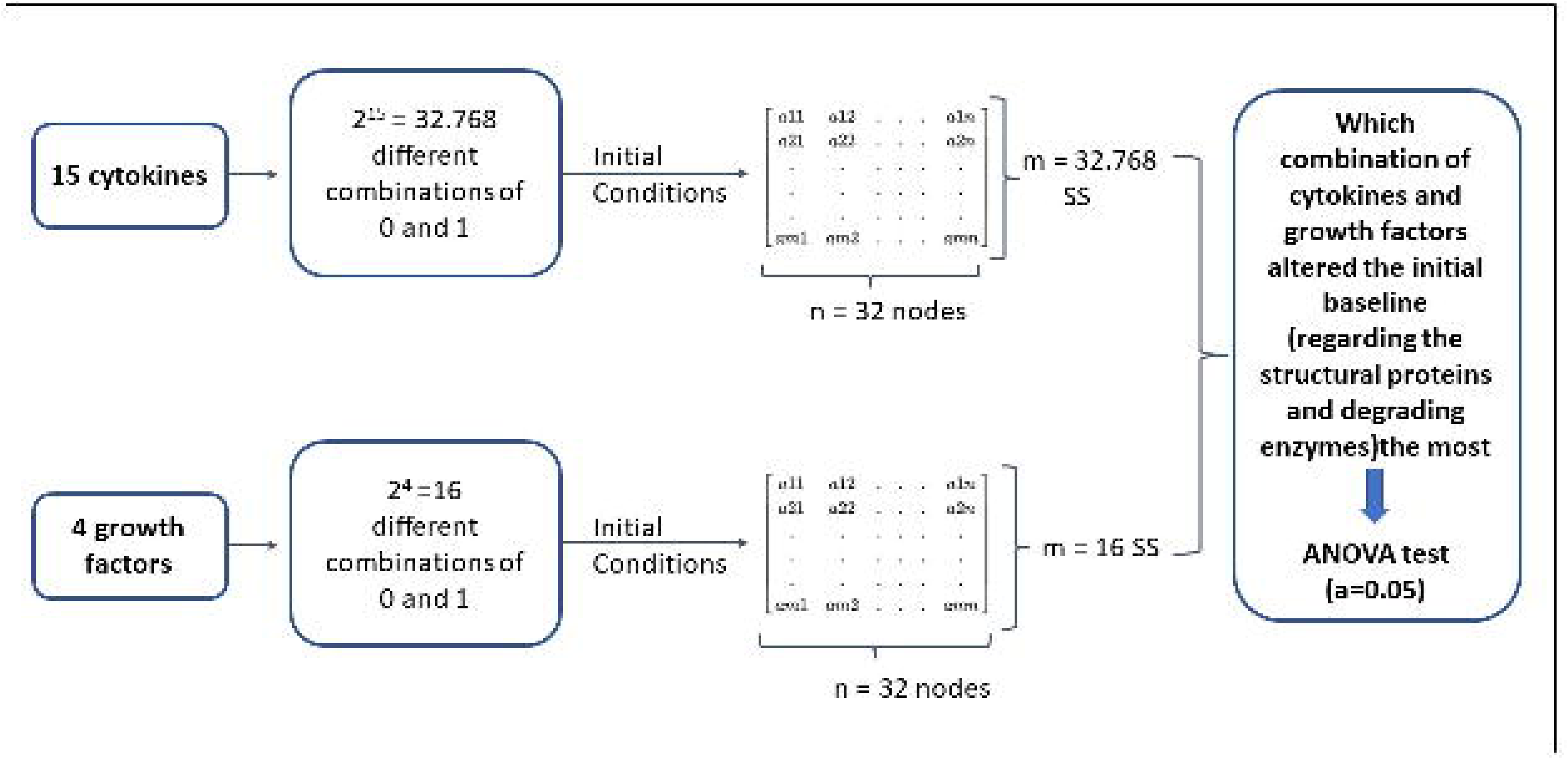
Schematic representation of the sensitivity analysis. As a first step of the sensitivity analysis a full factorial design of experiment was made to generate all the possible combinations among the 15 cytokines and the 4 growth factors, with two treatment levels (0 and 1). These combinations were then used as initial conditions to solve the system semi-quantitatively and finally get the corresponding SS for each combination. Finally, by using an ANOVA test, the combinations that altered the initial baseline the most was identified.

As a first step, a full factorial design of experiment was made and described by n^p^ total interactions, when n is the number of levels (0 and 1 in our case) and p is the number of factors/proteins tested. In that way all the possible combinations among the analysed proteins are generated and represent the initial states. More specifically, 2^15^= 32.768 simulated combinations among the 15 cytokines and 2^4^=16 simulated combinations among the growth factors were tested. For each combination, the direct effect and the second-order effect of the tested factors to the interest groups was investigated. The second step was to solve again our system semi-quantitatively so many times as the total combinations of each tested group, by using each combination as initial state, to obtain the relevant SS. This led to 32.768 different SS for the cytokines and 16 different SS for the growth factors.

The last step included the analysis of the experiments by using the analysis of variance F statistic (ANOVA F Statistic). Such a method compares the variability between the groups to the variability within the groups. The formula used for the ANOVA F Statistic for each node of the network-based model is:

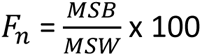

where *MSB* is the mean sum of squares between the groups

`o` cytokines-structural proteins
`o` cytokines-degrading enzymes
`o` growth factors-structural proteins
`o` growth factors-degrading enzymes

and *MSW* is the mean sum of squares within the groups (within the 15 cytokines and within the 4 growth factors). Thereof, the larger the F value, the greater is the evidence that there is a difference between the group means. To compare if the difference among the group means is statistically different, we computed the p-value. The significant F values (p ≤ 0.05) for the interested groups were plotted in Pareto charts in a logarithmic scale as the distribution for the proteins was very large. In the results only the direct effects of the cytokines can be seen for the sake of simplicity, but the analysis included the second-order interactions. Regarding the growth factors all the effects (direct and second order) can be visualized as they were few.

## Acknowledgements

Financial support was received from the Marie Skłodowska Curie International Training Network (ITN)“disc4all”(https://disc4all.upf.edu, accessed on 10 May 2022) grant agreement #955735 (https://cordis.europa.eu/project/id/955735, accessed on 10 December 2022)

## Author Contributions

S.T. wrote the main manuscript. S.T., M.S. and J.N. developed methods. ST, CLM, JP and JN provided intellectual input on the development of the network model, S.T. prepared the figures. All authors reviewed and approved the manuscript.

## Supplementary Material

The Supplementary Material can be found attached along with the manuscript.

**Competing Interests statements**

## Abbreviations

IDD: Intervertebral disc degeneration
ECM: Extracellular matrix
RNM: Regulatory network model
NPC: Nucleus pulposus cells
IVD: Intervertebral disc
TGF-β: Transforming growth factor beta
IL-1Ra: Interleukin-1 receptor antagonist
LBP: Low back pain
NPC: Nucleus pulposus cells
COL2A: Type-II collagen ACAN Aggrecan
MMP: Matrix metalloproteinases
ADAMTs: A disintegrin and metalloproteinase with thrombospondin motifs
IL-10: Interleukin 10
IL-4: Interleukin 4
GDF5: Growth differentiation factor 5
IGF1: Insulin-like growth factor 1
TIMP: Tissue Inhibitor of metalloproteinases
SS: Steady state
COL10A1: Collagen Type X Alpha 1
COL1A: Type-I collagen
IFN-γ: Interferon gamma
TNF: Tumor necrosis factor
IL-1α: Interleukin-1 alpha
IL-6: Interleukin 6
CCL C-C: Motif Chemokine Ligand
VEGF: Vascular endothelial growth factor

