## Supplementary Material for "Nucleus Pulposus Cell Network Modelling in the Intervertebral Disc"

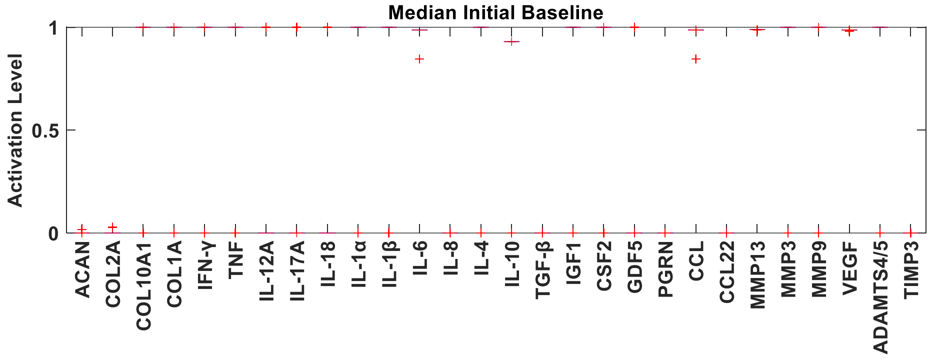


**Fig.1**. Boxplots of the initial RNM network. The baseline of the initial regulatory network model (RNM) for every protein is represented by red boxplots.

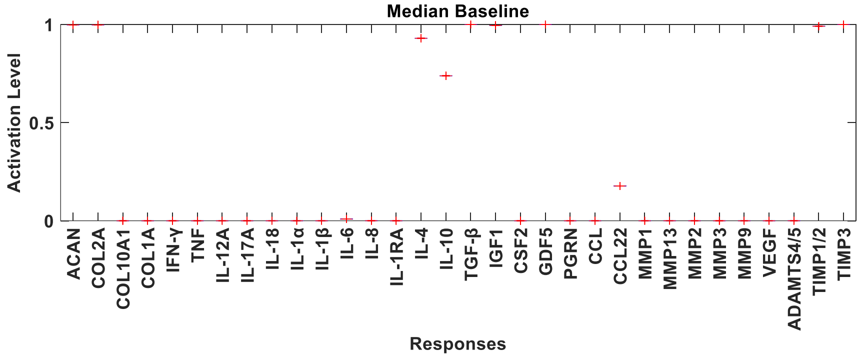


**Fig.2**. Boxplots of the initial RNM network. The baseline of the enriched regulatory network model (RNM) for every protein is represented by red boxplots.

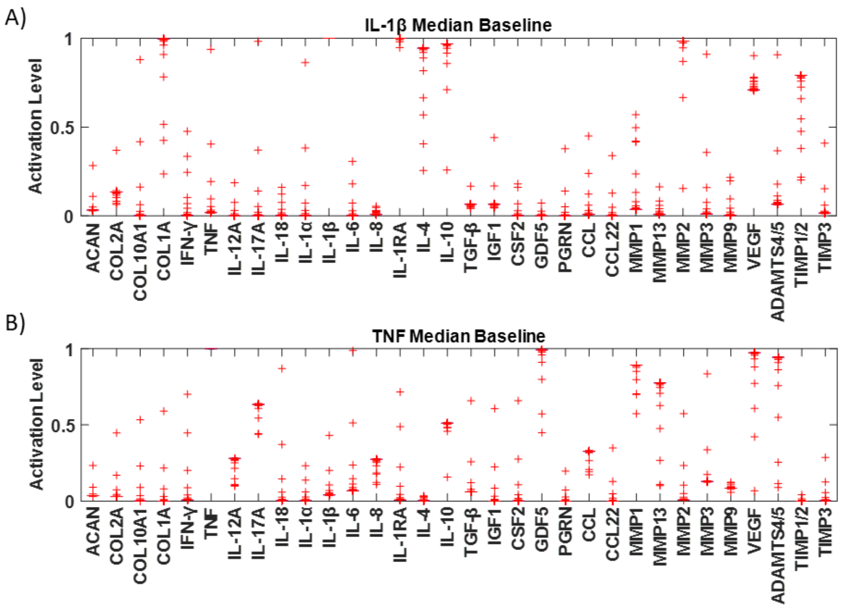


**Fig.3**. Boxplots of the perturbed baselines. The baseline of the enriched regulatory network model (RNM) for every protein after A) IL-1β stimulation and B) TNF stimulation is represented by red boxplots.

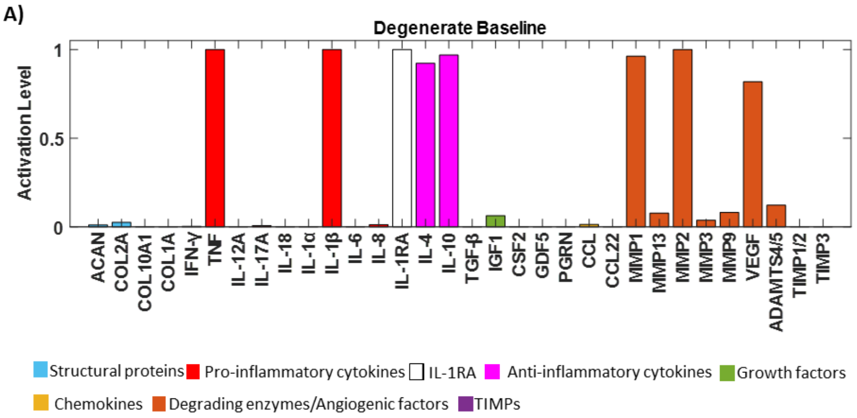

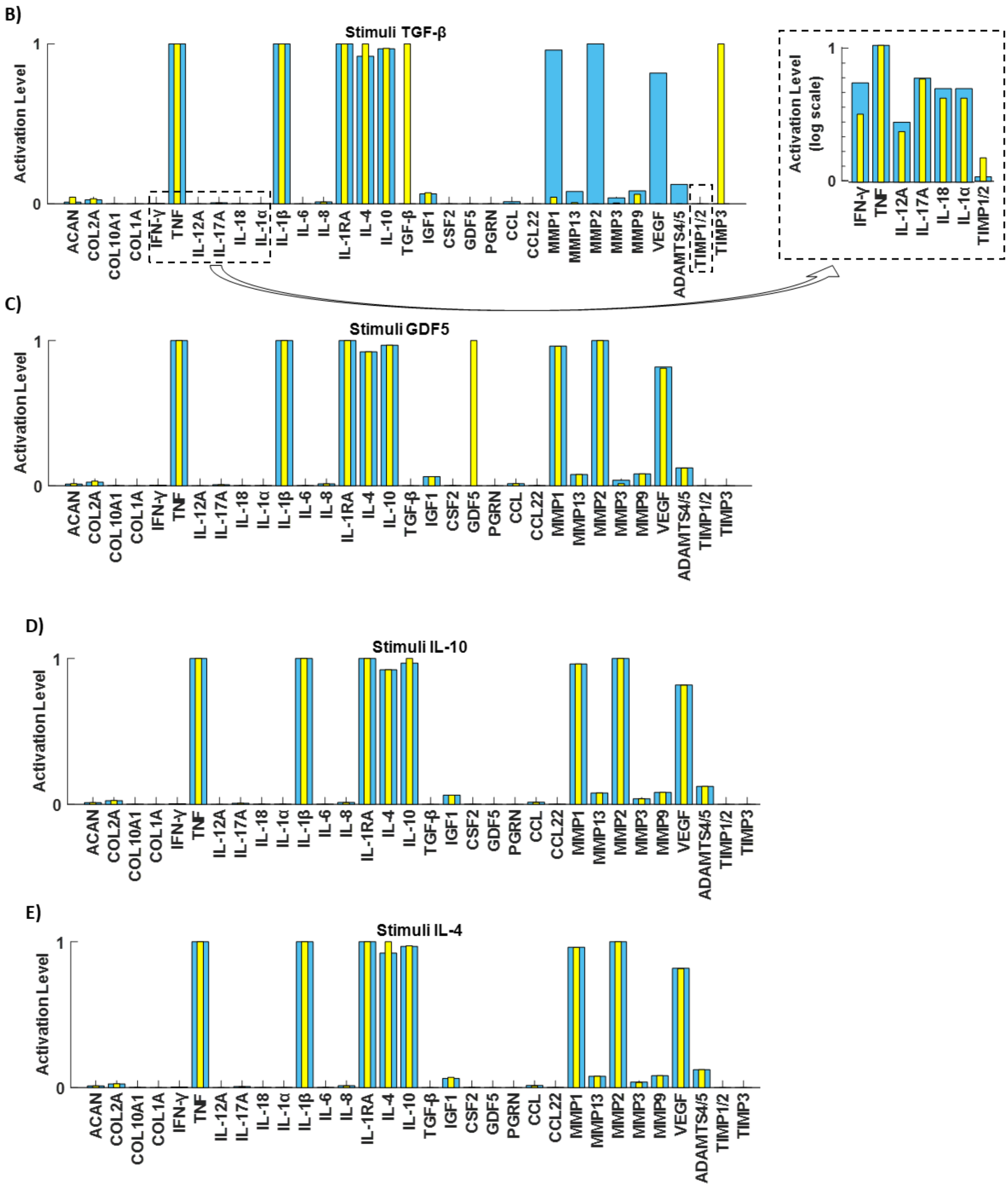


**Fig.4**. Assessment of the network through independent tests. A degenerate baseline was produced to promote catabolism by clamping TNF and IL-1β (Fig.3A) to 1. Rescue strategies were simulated by stimulating the degenerate baseline with B) TGF-β, C) GDF5, D) IL-10 and E) IL-4 (yellow bars).
